# β-Catenin Stabilization Protects Against Pulmonary Hemorrhage Through Amphiregulin and BATF- Mediated Regulatory T Cells

**DOI:** 10.1101/2025.10.21.683678

**Authors:** Fiona Mason, Hui Xiong, Ali Mobeen, Saddam Hossain, Sara Mahmudlu, Rosanne Trevail, Mikyal Mobeen, Li Chen, Sunny Lee, Tuncay Delibasi, Jyoti Misra Sen, Melanie Comito, Mobin Karimi

**Affiliations:** Department of Microbiology and Immunology, SUNY Upstate Medical University, Syracuse, NY 13210; School of Medical Imaging, Nanchang Medical College, China; Endocrinology, Diabetes and Metabolism Internal Medicine Medical Genetic SUNY Upstate Medical University, Syracuse, NY 13210; National Institute on Aging, NIH; Center on Aging and Immune Remodeling Immunology Program Department of Medicine Johns Hopkins University Baltimore, MD 21224; Division of Pediatric Hematology/Oncology Paige Yeomans Arnold Endowed Professor in Pediatric Oncology Upstate Medical University

**Keywords:** β-Catenin Stabilization, Pulmonary Hemorrhage, Amphiregulin, Transcriptional Factor BATF, Tissue-resident regulatory T cells

## Abstract

Pulmonary hemorrhaging (PH) is a life-threatening condition with a high mortality rate, yet the role of immune cells in its pathogenesis remains poorly defined. Here, we investigated the protective function of β-catenin stabilization in T cells and its impact on PH. Using a novel transgenic mouse model (CAT-Tg) with stabilized β-catenin, we demonstrate that β-catenin stabilization induces a distinct T-cell phenotype characterized by an expansion of central effector memory cells (CD44⁺, CD122⁺, Eomes⁺, T-bet⁺). Mechanistically, this effect was associated with suppression of key proinflammatory pathways, including reduced phosphorylation of STAT1, STAT3, and JAK1.

PH was induced using pristane, and CAT-Tg mice were significantly protected from lung damage, showing reduced proteinuria and decreased pulmonary proinflammatory cytokine production compared with wild-type (WT) and T cell–specific β-catenin knockout (cKO) mice. This protection correlated with a marked increase in FOXP3⁺ regulatory T cells (Tregs) in CAT-Tg mice. We further identified a novel mechanism in which β-catenin stabilization enhances lung expression of Amphiregulin and BATF, two molecules essential for Treg function and tissue repair. Adoptive transfer of CAT-Tg Tregs into WT mice with pristane-induced PH conferred superior protection, as evidenced by reduced lung inflammation and proteinuria. The systemic administration of a β-catenin agonist to mice with PH significantly attenuated disease severity. Our bioinformatic analysis confirmed that β-catenin stabilization upregulates pathways associated with tissue repair and immune homeostasis, including PI3K-Akt, angiogenesis, and STAT5 signaling. Collectively, these findings reveal that β-catenin stabilization protects against pulmonary hemorrhage by inducing a specialized T-cell phenotype and establishing a protective Amphiregulin–BATF–Treg axis. This study identifies a novel immunomodulatory pathway with therapeutic potential for PH and other inflammatory lung diseases.

## Introduction

Pulmonary hemorrhage (PH) is a severe and often fatal condition associated with autoimmune disorders such as Goodpasture syndrome, systemic lupus erythematosus (SLE), and antiphospholipid syndrome, in which autoimmune injury to alveolar capillaries causes bleeding into the lungs (1-6). PH also arises in diverse respiratory diseases including bronchiectasis, tumors, tuberculosis, aspergilloma, and cystic fibrosis, where it frequently presents as hemoptysis (7-10). Current treatments are largely supportive, focusing on ventilatory support, hemodynamic stabilization, and oxygenation (11). Pharmacological interventions such as antifibrinolytic agents promote clotting (3, 12-16), while immunosuppressive therapies aim to control inflammation (17-19). Despite these strategies, mortality remains high, and the cellular and molecular mechanisms by which immune cells exacerbate or mitigate PH are poorly understood. Addressing this critical gap is the focus of the present study.

Amphiregulin (AREG) and the transcription factor BATF are central mediators of tissue repair (20-26). Both regulate immune responses and facilitate regeneration of damaged tissue. β-catenin is a critical regulator of AREG expression as the β-catenin signaling pathway centrally controls the production of AREG (27, 28). β-Catenin signaling pathways play a critical role in T-cell development and tissue homeostasis (29). When activated, β-catenin accumulates in the cytoplasm, translocates to the nucleus, and interacts with T-cell factor/lymphoid enhancer factor (Tcf/Lef) transcription factors (30-32). This complex binds to promoter regions of target genes to initiate transcription, including direct induction of AREG (27, 33). Although β-catenin regulates AREG expression, no direct molecular link between β-catenin and BATF has been established. β-Catenin is also known to promote transcriptional programs involved in proliferation, migration, and stem cell maintenance during injury repair (29, 34). In parallel, BATF, a component of the AP-1 transcription factor complex, is essential for differentiation and function of multiple immune subsets, including T cells (35) and innate lymphoid cells (ILCs) (36). Another key regulator of immune tolerance and tissue repair is the FOXP3⁺ regulatory T cell (Treg) population. AREG has been shown to enhance Treg suppressive activity (21, 37). Yet, the upstream molecular pathways linking β-catenin, BATF, and AREG in Tregs remain undefined.

Here, we demonstrate that β-catenin regulates FOXP3 expression and enhances BATF and AREG function in Tregs, thereby protecting against PH. Using transgenic mice with β-catenin stabilization (CAT-Tg) (38-40) and conditional T cell–specific β-catenin knockout (cKO), we demonstrate that CAT-Tg CD8⁺ T cells display increased central and effector memory subsets, elevated expression of CD44, CD122, Eomes, and T-bet, and reduced phosphorylation of STAT1, STAT3, and JAK1. Functionally, CAT-Tg mice were protected from pristane-induced PH, exhibiting reduced lung pathology, less proteinuria, decreased proinflammatory cytokines, and enhanced anti-inflammatory responses compared with WT and β-catenin cKO control mice.

Importantly, CAT-Tg mice showed increased frequencies of Tregs expressing high levels of AREG and BATF. Adoptive transfer of CAT-Tg Tregs into WT recipients with pristane-induced PH conferred significant protection, reducing lung inflammation and proinflammatory cytokines while enhancing IL-10 production. Pharmacologic activation of β-catenin in WT mice similarly protected against PH, reducing inflammatory cytokines and modestly expanding Tregs. Finally, unbiased RNA sequencing revealed that β-catenin stabilization reprograms gene expression toward tissue repair and immune homeostasis.

Collectively, these findings identify a previously unrecognized β-catenin–AREG–BATF–Treg axis that protects against pulmonary hemorrhage. This work not only uncovers a novel immunoregulatory pathway but also highlights β-catenin as a potential therapeutic target for PH and other cytokine storm–mediated lung disease.

## Results

### CAT-Tg mice selectively impact the CD8⁺ T-cell phenotype

β-Catenin is required for both αβ and γδ T cell development (41-43). To determine whether stabilization of β-catenin under the proximal Lck promoter alters specific αβ T-cell populations, we analyzed T-cell subsets from wild-type (WT) and CAT-Tg mice. The αβ T-cell phenotype is critical because it reflects differentiation status, effector function, and potential for persistence and therapeutic efficacy (44-47). Freshly isolated splenic CD3⁺ T cells were stained with CD44 and CD62L and analyzed by flow cytometry. Gating was performed on total T cells, which were then separated into CD4⁺ and CD8⁺ populations. Each subset was further evaluated for naïve, central memory (CM), and effector memory (EM) cells based on CD44 and CD62L expression (38, 48-52). CD62L (L-selectin) facilitates homing to lymphoid tissues, whereas CD44 is associated with activation, adhesion, and migration to peripheral tissues or inflammatory sites (53-56). Our analysis revealed that CD8⁺ T cells from CAT-Tg mice exhibited a significantly higher proportion of CM and EM subsets compared with WT controls. Data with gating is shown in representative flow plots along with quantitative analysis of the data (Fig. 1A–E). In contrast, CD4⁺ T cells showed no significant differences in the distribution of naïve, CM, or EM populations between CAT-Tg and WT mice (Fig. 1F-J). These findings indicate that β-catenin stabilization preferentially drives memory differentiation in CD8⁺ T cells, a subset critical for long-term antitumor immunity and therapeutic efficacy.

**Figure 1.**
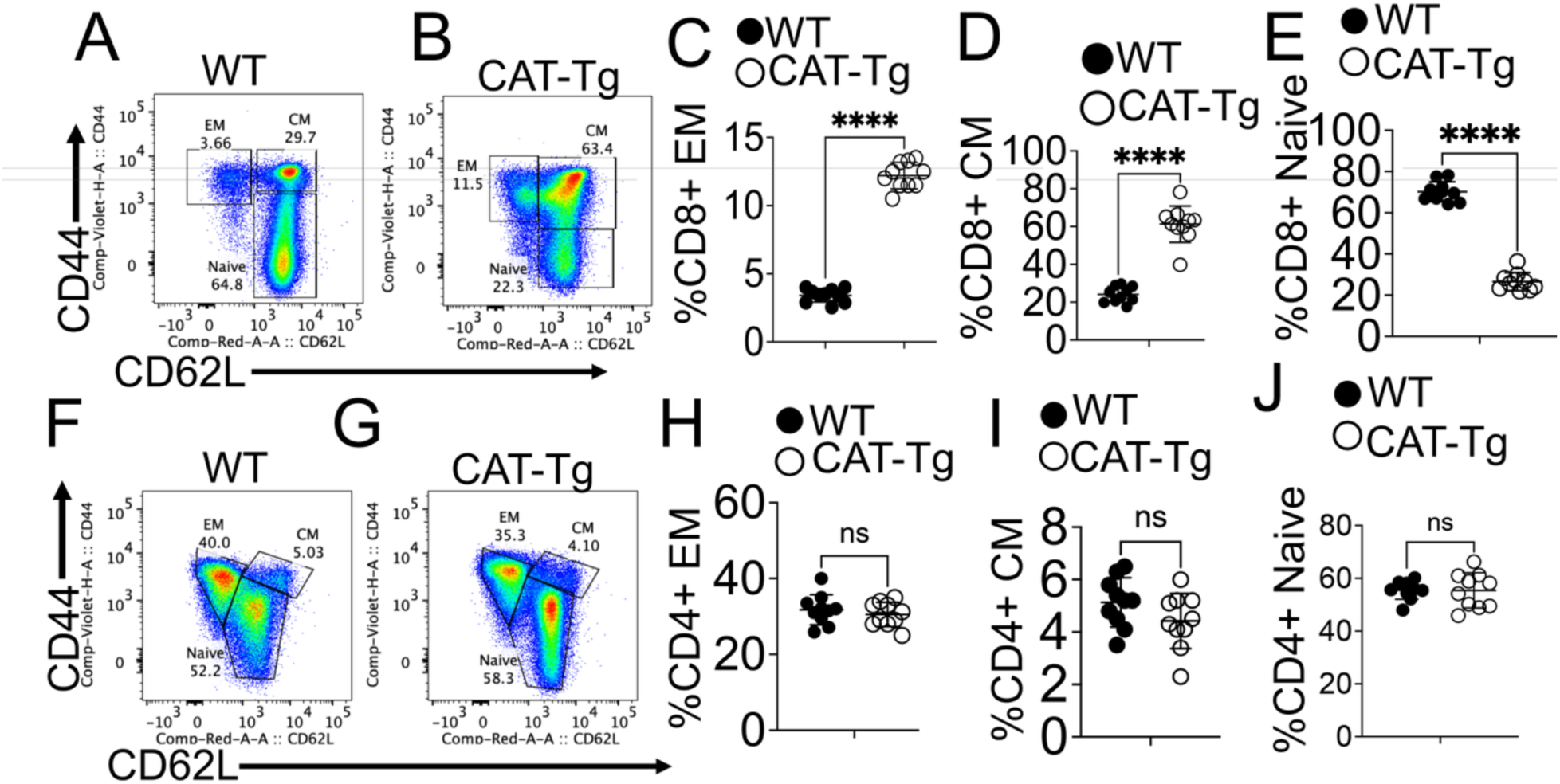
CAT-Tg mice selectively alter the CD8⁺ T-cell phenotype: Splenocytes from WT and CAT-Tg mice were analyzed for central memory (CM) and effector memory (EM) phenotypes. Freshly isolated spleen cells were first gated on CD3⁺ T cells, and then separated into CD4⁺ and CD8⁺ subsets. Within the CD8⁺ population, expression of CD44 and CD62L was examined by flow cytometry (A–B). CD44⁺CD62L⁻ cells were classified as EM T cells, CD44⁺CD62L⁺ cells as CM T cells, and double-negative cells as naïve CD8⁺ T cells. Representative flow cytometry plots from WT and CAT-Tg mice illustrate these subsets, with quantitative comparisons shown for EM, CM, and naïve CD8⁺ T cells (C–D). Similarly, CD4⁺ T cells were analyzed using the same gating strategy. Representative plots (F–G) and quantitative analyses demonstrate the distribution of EM, CM, and naïve subsets within CD4⁺ T cells. All quantitative data are presented as mean ± standard error of the mean (SEM). Sample sizes (*n* = 10 mice per group) are indicated in the figure panels. Statistical significance was determined using the appropriate test, with ****p < 0.0001.

### CD8⁺ T cells from CAT-Tg mice express higher activation markers but reduced phosphorylation of key signaling molecules

We previously reported that T cells with attenuated TCR signaling display increased activation markers without inducing alloimmunity (50, 51, 57-59). To test whether this was also true in CAT-Tg mice, we examined CD44 expression on CD8⁺ T cells. CD44 serves as a marker of activation and memory, while also facilitating T-cell migration, tissue infiltration, adhesion, and survival during immune responses (60). Flow cytometric analysis revealed a significant increase in CD44 expression on CD3⁺CD8⁺ T cells from CAT-Tg mice compared with WT controls, as shown in representative plots and quantitative analyses (Fig. 2A–B). We next assessed transcription factors that regulate CD8⁺ T-cell memory. T-bet and Eomes are key T-box factors that cooperate to drive memory formation by promoting CD122 expression, which is essential for IL-2 and IL-15 signaling (61, 62). Consistent with enhanced memory potential, CD8⁺ T cells from CAT-Tg mice exhibited significantly higher levels of T-bet, Eomes, and CD122 compared with WT CD8⁺ T cells (Fig. 2C–H).

**Figure 2:**
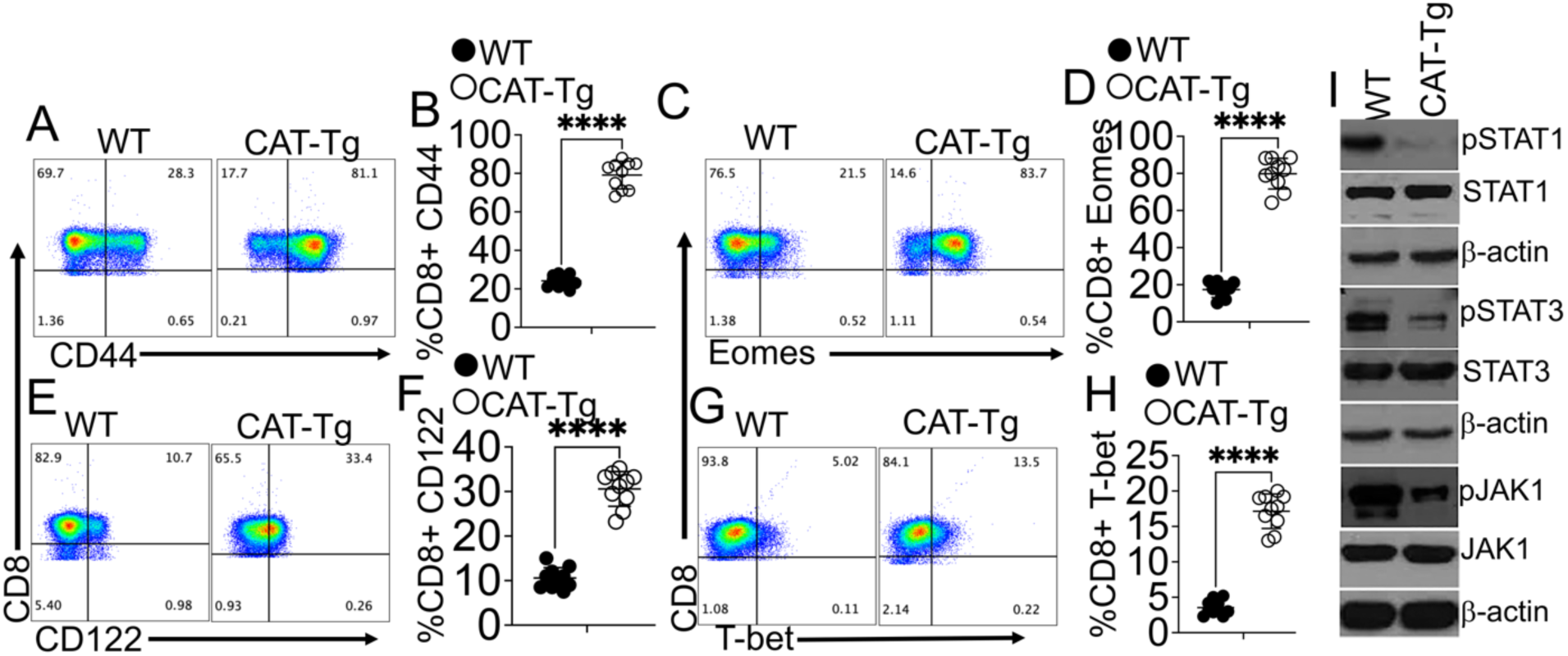
CD8⁺ T cells from CAT-Tg mice express higher activation markers but reduced phosphorylation of key signaling molecules: Freshly isolated spleen cells were gated on CD3⁺ and CD8⁺ T cells from WT and CAT-Tg mice. These CD3⁺CD8⁺ T cells were further gated for the CD44 activation marker, as shown by flow-cytometry plots (A). Flow-plot data from 10 WT and 10 CAT-Tg mice were quantified and presented using Prism plots (B). CD8⁺ T cells from WT or CAT-Tg mice were assessed for CD122 (flow plots, E) with quantitative analysis of 10 WT and 10 CAT-Tg mice (F). Next, the same CD8⁺ T cells were stained intracellularly for Eomes and T-bet, as shown by flow plots (C–G), with quantitative analysis of 10 WT and 10 CAT-Tg mice. CD3⁺ T cells from each mouse were MACS-purified; 1 × 10⁷ T cells were pulsed for 3 minutes with CD3/CD28 for activation and then lysed. Cell lysates were examined for phosphorylated STAT1 (pSTAT1), total (non-phosphorylated) STAT1, and β-actin as a control. Cells were also examined for pSTAT3, total (non-phosphorylated) STAT3, JAK1, and β-actin as a control. All quantitative data are presented as mean ± standard error of the mean (SEM). Sample sizes (n = 10 mice per group) are indicated in the figure panels. Statistical significance was determined using the appropriate test, with ****p < 0.00

Finally, we examined whether stabilization of β-catenin alters proximal T-cell signaling. STAT1, STAT3, and JAK1 are central mediators of the JAK-STAT pathway, controlling T-cell growth, differentiation, and immune responses (63, 64). Upon CD3/CD28 stimulation, CAT-Tg T cells displayed significantly reduced phosphorylation of STAT1, STAT3, and JAK1 compared with WT controls (Fig. 2I). These findings suggest that β-catenin stabilization promotes a highly activated and memory-prone CD8⁺ T-cell phenotype, while simultaneously dampening canonical JAK-STAT signaling.

### β-catenin stabilization protects mice from pulmonary hemorrhage (PH)

To determine whether β-catenin stabilization exacerbates or ameliorates pulmonary hemorrhage (PH), we utilized 6–8-week-old WT, CAT-Tg, and β-catenin cKO mice, all on a C57BL/6 (B6) background. Pre-treatment urine and serum samples were collected, then the mice were injected intraperitoneally with 0.5 mL/20 g of Pristane (44, 65, 66). Mice from each group were monitored for weight loss and clinical signs of disease. No significant weight loss was observed within 14 days. On day 14, mice were euthanized, and lungs were collected (Fig. 3A). Gross examination revealed that lungs from CAT-Tg mice were markedly protected from PH compared with those from WT and β-catenin cKO mice (Fig. 3B–D). Histological analysis of lung tissue sections stained with hematoxylin and eosin (H&E) further confirmed reduced hemorrhage in CAT-Tg mice compared with those from WT and β-catenin cKO mice. Importantly, all lung sections were independently evaluated by a blinded pathologist (Li Chen), and double-blind pathological scoring validated the protective effect of β-catenin stabilization (Fig. 3E). We also examined other organs including kidneys, liver and spleen. We did not see any significant differences among WT, CAT-Tg and β-catenin cKO mice within 14 days (Sup. Fig. 1A-O). However, we saw significant differences in kidneys, livers, and spleen over a 3 month time-period among these mice (data not shown).

**Figure 3.**
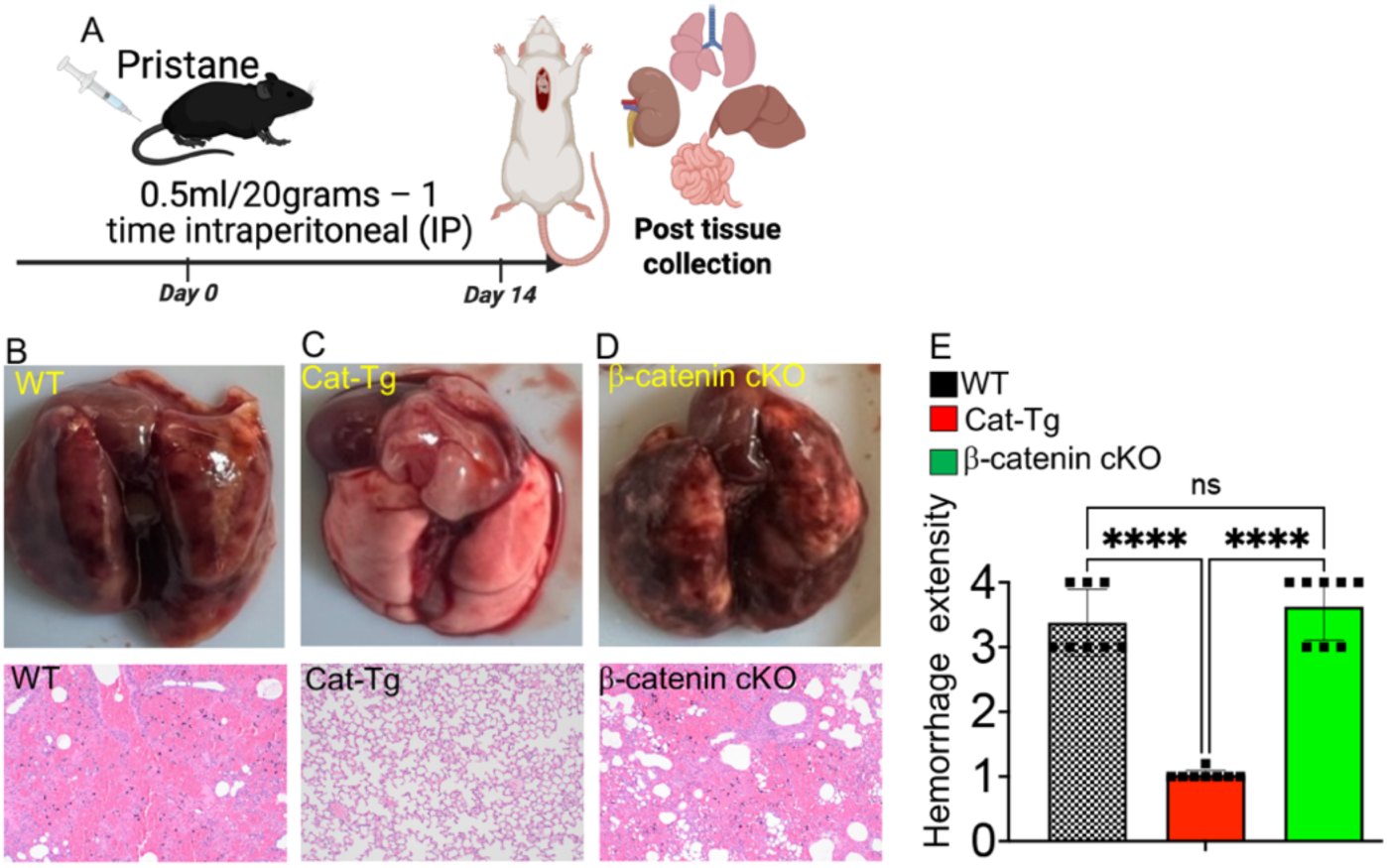
β-catenin stabilization protects mouse lungs from pulmonary hemorrhage (PH). (A) Schematic representation of the experimental design. WT, CAT-Tg, and β-catenin cKO mice were intraperitoneally (IP) injected with 0.5 ml/g Pristane. On day 14, mice were euthanized, and lungs, liver, kidneys, and small intestine were harvested. (B–D) Representative gross anatomical images of lungs from WT, CAT-Tg, and β-catenin cKO mice (upper panels). Corresponding histological sections of lungs were stained with H&E (lower panels). (E) Histological slides were evaluated and scored by a pathologist. Quantitative data are shown as mean ± standard error of the mean (SEM). Sample sizes (n = 10 mice per group) are indicated in the figure panels. Statistical significance was assessed using the appropriate test, with ****p < 0.0001

These results demonstrate that β-catenin stabilization protects mice from pristane-induced PH. Having established that β-catenin stabilization confers protection against PH, we next sought to investigate the underlying immunological mechanisms driving this protective effect.

### β-Catenin stabilization reduces proteinuria, suppresses pro-inflammatory cytokines, and enhances anti-inflammatory cytokines during PH

Proteinuria, defined as excess protein in the urine, often accompanies pulmonary hemorrhage (PH) as part of a pulmonary–renal syndrome. This syndrome is most commonly caused by systemic autoimmune disorders that simultaneously damage both the kidneys and lungs, and early detection is critical to prevent organ failure (67-70). To evaluate kidney involvement, we measured proteinuria by ELISA in WT, CAT-Tg, and β-catenin cKO mice (Fig. 4A). WT and β-catenin cKO mice treated with pristane exhibited significantly increased proteinuria by day 14 compared with pretreatment levels. In contrast, CAT-Tg mice showed no difference in proteinuria before and after pristane treatment, providing evidence that β-catenin stabilization protects organs from PH-associated damage (Fig. 4B–D).

**Figure 4.**
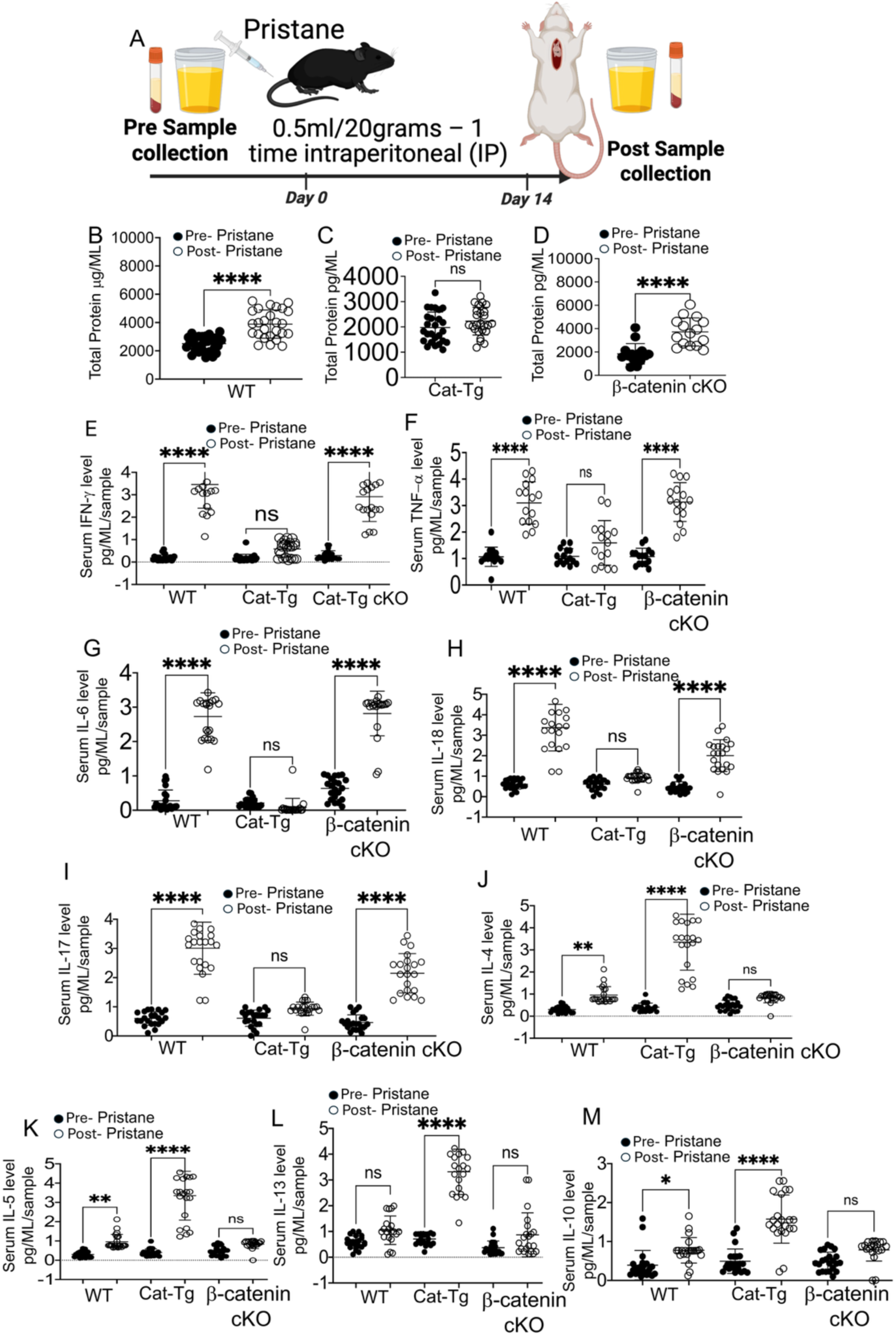
β-catenin stabilization reduces proteinuria, suppresses pro-inflammatory cytokines, and enhances anti-inflammatory cytokines during PH. (A) Schematic representation of the experimental design. WT, CAT-Tg, and β-catenin cKO mice were intraperitoneally (IP) injected with 0.5 ml/g Pristane. Urine and blood were collected via tail vein prior to injection and again on day 14, when mice were euthanized. (B–D) Proteinuria was measured by ELISA in each group, with quantitative comparison of pre- and post-Pristane samples. (E–M) Cytokine profiles were analyzed using LEGENDplex, including IFN-γ, TNF-α, IL-6, IL-18, IL-17, IL-4, IL-5, IL-13, and IL-10, and compared between pre- and post-Pristane conditions. Quantitative data are presented as mean ± standard error of the mean (SEM). Sample sizes (n = 15–25 mice per group) are indicated in the figure panels. Statistical significance was determined by one-way ANOVA, with **p < 0.01, ***p < 0.001, and ****p < 0.0001.

Next, we assessed inflammatory cytokines. Interferon-γ (IFN-γ) and tumor necrosis factor-α (TNF-α) are key drivers of PH during systemic inflammation (71-73). WT and β-catenin cKO mice exhibited significant increases in both cytokines at day 14 post-pristane injection compared with baseline, whereas CAT-Tg mice showed no change in IFN-γ or TNF-α between pre- and post-treatment (Fig. 4E–F). Because IL-6 and IL-18 are important pro-inflammatory mediators of lung injury and hemorrhage (74-78), we examined their serum levels. IL-6 recruits neutrophils, a hallmark of hemorrhagic inflammation, while IL-18 amplifies inflammasome-driven cascades leading to tissue damage, vascular permeability, and alveolar bleeding. CAT-Tg mice exhibited significantly lower levels of both IL-6 and IL-18 compared with WT and β-catenin cKO mice following PH induction (Fig. 4G–H). IL-17 has also been implicated in lung injury, edema, and hemorrhage (79-81). Consistent with a protective effect, CAT-Tg mice displayed significantly reduced IL-17 induction compared to WT and β-catenin cKO mice (Fig. 4I), suggesting that β-catenin stabilization suppresses multiple pro-inflammatory pathways to protect against PH. We then tested whether β-catenin stabilization enhances anti-inflammatory cytokines. Indeed, serum from CAT-Tg mice contained significantly higher levels of IL-4, IL-5, IL-13, and IL-10 compared to controls (Fig. 4J–M). In contrast, no significant differences were observed in IL-12 or IL-9 levels among the groups (Sup. Fig. 2), indicating that the protective effect was specific to certain anti-inflammatory cytokines (82, 83).

Together, these findings demonstrate that β-catenin stabilization protects against PH not only by suppressing pro-inflammatory cytokines such as IFN-γ, TNF-α, IL-6, IL-18, and IL-17, but also by enhancing anti-inflammatory cytokines including IL-4, IL-5, IL-13, and IL-10. Having established both structural and cytokine-level protection, we next investigated whether β-catenin-driven T cells exhibit functional advantages in vivo.

### β-Catenin stabilization enhances amphiregulin, and BATF expression in Tregs

We previously reported that regulatory T cells (Tregs) play a critical role in controlling alloimmunity (50, 51, 57-59, 84, 85). Tregs are also known to protect against PH by dampening excessive immune responses and promoting tissue repair (86, 87). To evaluate whether β-catenin stabilization impacts Tregs, we analyzed both spleens (Fig. 5 A-C) and lungs (Fig. 5 D-E) from WT, CAT-Tg and β-catenin cKO mice, gating by CD3⁺CD4⁺ T cells and determining CD25 and FOXP3 expression. CAT-Tg mice exhibited a significantly higher frequency of Tregs, including both conventional CD25⁺FOXP3⁺ and non-conventional CD25⁻FOXP3⁺ (84, 85) subsets, compared with WT and β-catenin cKO mice (Fig. 5A–C, F). These findings suggest that β-catenin stabilization protects the lung during PH, at least in part, by expanding the Treg population.

**Figure 5.**
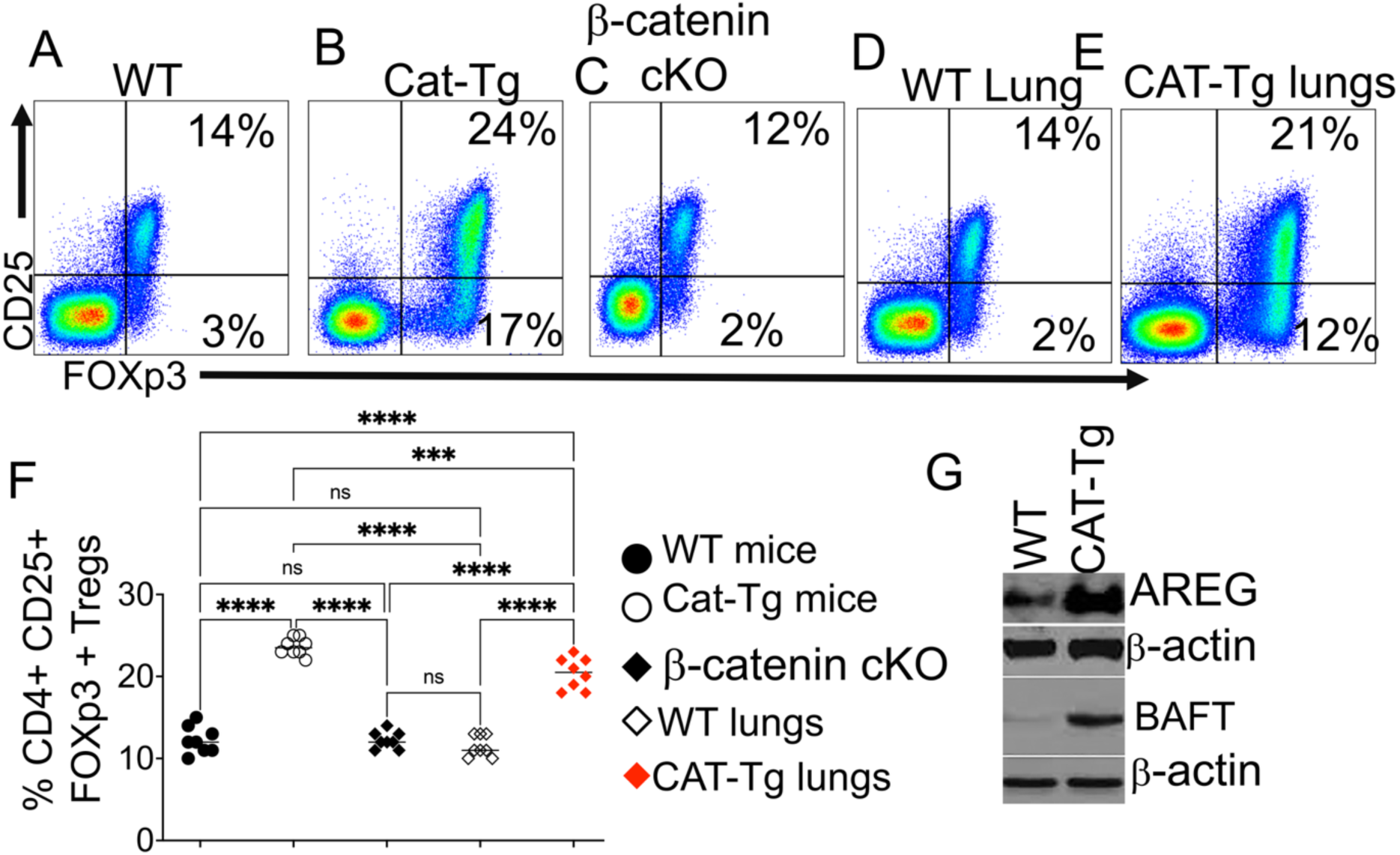
β-catenin stabilization enhances Tregs, amphiregulin, and BATF expression. (A–C) Flow cytometry analysis of freshly isolated splenocytes from WT, CAT-Tg, and β-catenin cKO mice. Cells were gated on CD3⁺, CD4⁺, and CD25⁺, followed by intracellular staining for the transcription factor FOXP3. (D–E) Lung tissues from WT and CAT-Tg mice were examined for Tregs using CD3⁺, CD4⁺, CD25⁺, and FOXP3 markers. (F) Representative flow cytometry plots showing gating on CD4⁺CD25⁺FOXP3⁺ Tregs, with quantification performed using Prism. (G) Tregs were FACS-sorted from lung tissue based on CD25⁺ and FOXP3 expression; protein lysates were immunoblotted for amphiregulin and BATF, with β-actin as the loading control. Quantitative data are presented as mean ± standard error of the mean (SEM). Sample sizes (n = 15–25 mice per group) are indicated in the figure panels. Statistical significance was determined by one-way ANOVA, with **p < 0.01, ***p < 0.001, and ****p < 0.0001.

To further define the protective mechanisms of Tregs, we examined amphiregulin (Areg), an epidermal growth factor receptor (EGFR) ligand that promotes tissue repair independent of Tregs’ classical immune-suppressive functions (37, 88-90). We FACS-sorted CD25⁺FOXP3⁺ Tregs from CAT-Tg and WT mice lungs by CD3^+^, CD4^+^, CD25^+^ and FOXp3^+^ (Fig. 5 D-E) by FOXp3 genetically tagged with Red fluorescent protein (RFP tag) and examine Areg expression. Our data revealed that Tregs from CAT-Tg mice expressed substantially higher levels of Areg, as demonstrated by Western blot analysis (Fig. 5G). These results highlight that β-catenin stabilization enhances both the abundance and tissue-repair capacity of Tregs.

Finally, we assessed expression of Basic leucine zipper ATF-like transcription factor (BATF), which is required for Treg homeostasis, differentiation, and stability (26). CAT-Tg Tregs expressed significantly higher BATF levels than WT controls, consistent with enhanced Treg stability (Fig. 5G). Because BATF sustains FOXP3 expression, this finding indicates that β-catenin stabilization not only expands Treg populations but also reinforces their lineage stability and suppressive function, thereby preventing uncontrolled inflammation and autoimmunity.

Together, these results demonstrate that β-catenin stabilization promotes a protective Treg program characterized by expansion, tissue-repair activity, and enhanced lineage stability. We next investigated whether these immunological changes translate into improved survival and disease outcomes in vivo.

### Adoptive transfer of CAT-Tg Tregs rescues pulmonary hemorrhage in vivo

To test whether Tregs from CAT-Tg mice, characterized by higher expression of Areg and BATF, confer in vivo protection against PH, we performed adoptive transfer experiments. WT mice were divided into three cohorts: untreated controls, recipients of WT Tregs, and recipients of CAT-Tg Tregs. All mice were injected intraperitoneally with pristane (0.5 mL/20 g body weight). Urine and serum were collected before treatment, and at day 10 post-injection splenic CD3⁺CD4⁺CD25⁺FOXP3⁺ Tregs were FACS-sorted from either WT or CAT-Tg donors. Each recipient received 1×10⁶ Tregs by adoptive transfer, while one cohort remained untreated. At day 21 post-pristane injection, urine, serum, and major organs (lungs, liver, intestine, kidney) were harvested (Fig. 6A–D). Gross pathology revealed that lungs from untreated pristane-injected WT mice displayed severe hemorrhagic damage (Fig. 6D). In contrast, mice rescued with CAT-Tg Tregs showed complete protection, with lungs appearing normal. Mice receiving WT Tregs exhibited partial protection, with reduced but still visible damage compared with untreated controls. Consistent with the histological findings, proteinuria was markedly elevated in untreated pristane-injected WT mice compared with pre-treatment levels. Adoptive transfer of CAT-Tg Tregs completely prevented proteinuria, whereas WT Tregs only partially reduced it (Fig. 6E). These data demonstrate that CAT-Tg Tregs more effectively protect renal function during PH.

**Figure 6.**
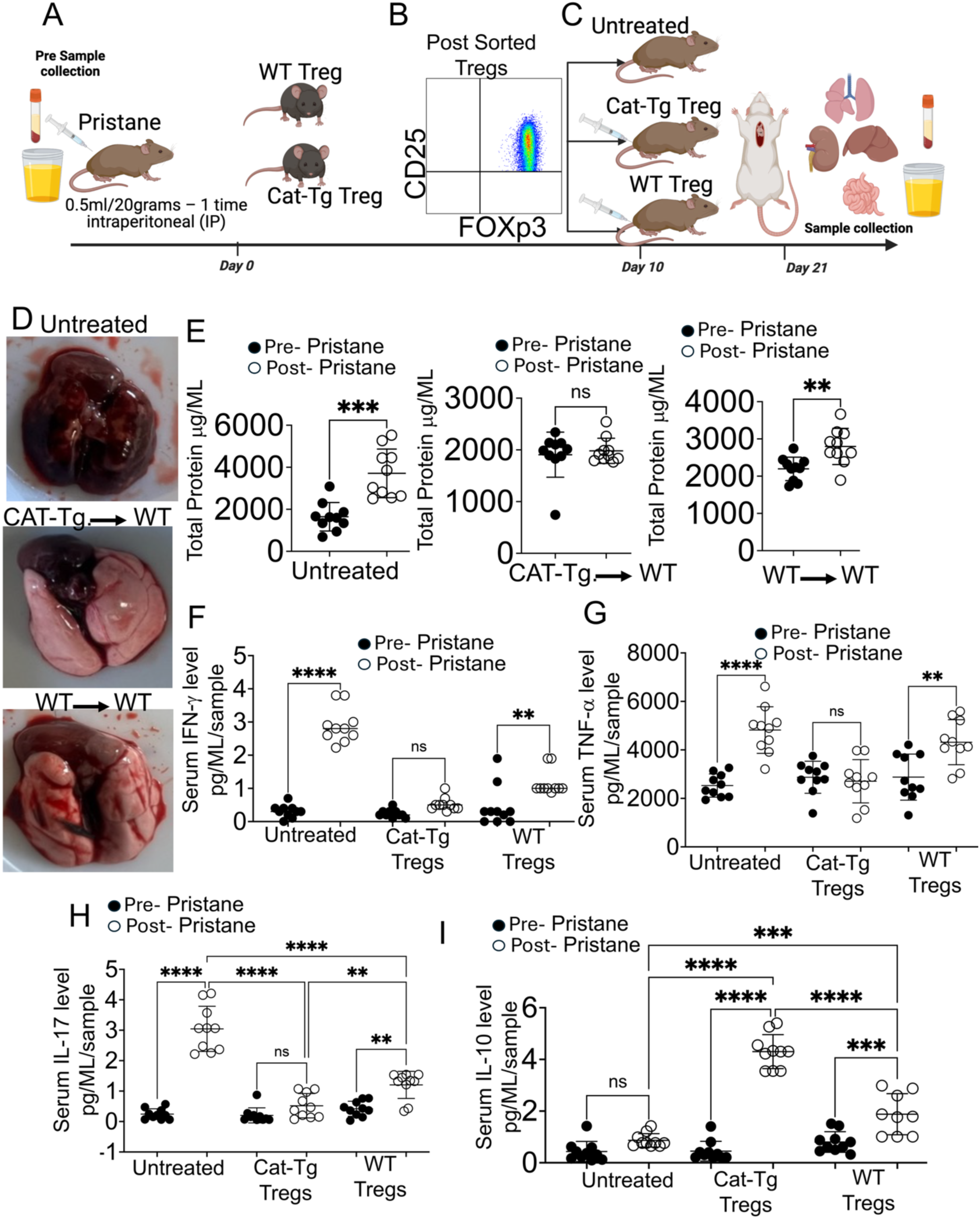
Adoptive transfer of CAT-Tg Tregs rescues pulmonary hemorrhage in vivo. (A) Schematic representation of the experimental design. WT mice were divided into three groups, and urine and blood were collected via tail vein. (B) Tregs were isolated from WT and CAT-Tg mice based on CD25⁺ and FOXP3 expression. (C) One cohort of mice was left untreated, a second cohort received 1 × 10⁶ Tregs from WT mice, and a third cohort received 1 × 10⁶ Tregs from CAT-Tg mice. On day 21, host mice were euthanized, and lungs, liver, kidneys, and small intestine were harvested. (D) Representative lung images from untreated Pristane-injected WT mice, WT mice treated with Tregs from WT donors, and WT mice treated with Tregs from CAT-Tg donors. (E) Proteinuria was measured by ELISA in each group, comparing pre- and post-Pristane samples. (F) Serum levels of IFN-γ, TNF-α, IL-17, and IL-10 were quantified in untreated mice and in mice treated with Tregs from WT or CAT-Tg donors, both pre- and post-Pristane injection. Quantitative data are presented as mean ± standard error of the mean (SEM). Sample sizes (n = 15–25 mice per group) are indicated in the figure panels. Statistical significance was determined by one-way ANOVA, with **p < 0.01, ***p < 0.001, and ****p < 0.0001.

Cytokine analysis further confirmed differential protective effects. WT mice rescued with CAT-Tg Tregs exhibited significantly reduced IFN-γ levels compared with those receiving WT Tregs (TNF-α) (Fig. 6F). IL-17 levels remained unchanged in CAT-Tg Treg recipients, whereas WT Tregs only modestly reduced IL-17 compared with untreated mice, and still showed higher levels relative to CAT-Tg Treg recipients (Fig. 6H). Importantly, CAT-Tg Tregs also induced a robust increase in IL-10, a key anti-inflammatory cytokine, compared with both untreated and WT Treg-treated mice (Fig. 6I). Together, these results provide strong evidence that β-catenin-stabilized Tregs protect against PH by suppressing pro-inflammatory cytokines (IFN-γ, IL-17) while enhancing IL-10 production.

Finally, to confirm the persistence and trafficking of donor Tregs, we repeated adoptive transfers using congenic CD45.2 WT recipients and CD45.1 donor mice. At day 21 post-transfer, both donor- and host-derived Tregs were detected in the spleen (Sup. Fig. 3A–C). Immunohistochemistry of lung tissue revealed donor Tregs within inflamed lungs, confirmed by staining for CD4 (green), donor CD45.2 (blue), and FOXP3 (red), with merged images demonstrating colocalization (Sup. Fig. 3D–J). These findings confirmed successful engraftment and lung homing of donor CAT-Tg Tregs.

These results demonstrate that adoptively transferred CAT-Tg Tregs, enriched for Areg and BATF, confer superior protection against PH compared to WT Tregs by reducing pro-inflammatory cytokines, enhancing IL-10 production, and directly trafficking to lung tissue.

### β-Catenin agonists recapitulate the protective effects of genetic stabilization

To determine whether a pharmacological approach could mimic genetic β-catenin stabilization in a mouse model, we used several Wnt/β-catenin agonists from MedChem Express including (MCE cat# HY-114321) (91-93). WT mice were injected with pristane as described above, and urine and blood were collected for serum analysis. A cohort of mice received either vehicle or the β-catenin agonist. Each mouse was treated with 10 µg/20 gram of mouse mice of the drug twice a week for 14 days (Fig. 7A–B). On day 14, the animals were euthanized, and their lungs were examined for Treg expression. Our data confirmed that mice treated with β-catenin agonists showed an increased percentage of Tregs (Fig. 7C). We then examined lungs from mice that were either untreated, treated with pristane, or treated with either vehicle or β-catenin agonists. Mice treated with pristane, or vehicle showed significant lung damage compared with mice not injected with pristane. Strikingly, mice injected with pristane and treated with β-catenin agonists showed no signs of lung damage, as confirmed by gross pathology, H&E staining, and independent pathologist analysis (Fig. 7D–J).

**Figure 7.**
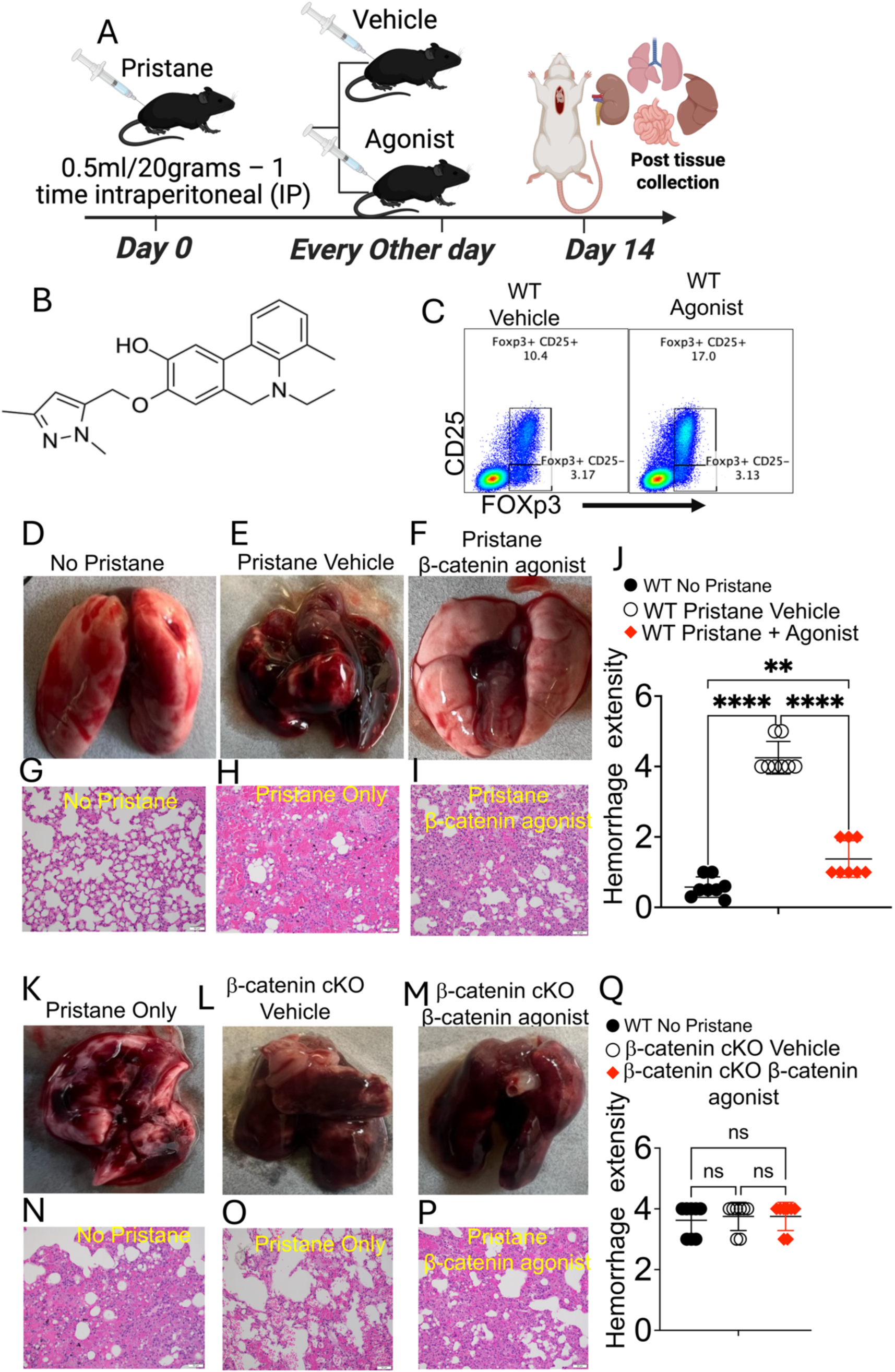
β-catenin agonists recapitulate the protective effects of genetic stabilization. (A) Schematic representation of the experimental design. WT mice were intraperitoneally injected with 0.5 ml/g Pristane and divided into two groups: one treated with vehicle and the other treated with a β-catenin agonist twice per week. On day 14, mice were euthanized, and lungs, liver, kidneys, and small intestine were collected. (B) Chemical structure of the β-catenin agonist. (C) Representative flow cytometry analysis of lung Tregs (CD4⁺CD25⁺FOXP3⁺) from WT mice treated with vehicle or β-catenin agonist. (D) Lung images from untreated mice that did not receive Pristane and had no detectable Tregs. (E–F) Lungs from Pristane-injected WT mice treated with either vehicle or β-catenin agonist. (G–J) Histological analysis (H&E staining) and scoring of lung tissues shown in panels D–F. (K) Lungs from Pristane-injected WT mice that received no vehicle or agonist treatment. (L–M) Lungs from β-catenin cKO mice treated with either vehicle or β-catenin agonist. (N–Q) Histological analysis and scoring of lung tissues from panels K–M. Quantitative data are presented as mean ± standard error of the mean (SEM). Sample sizes (n = 15–25 mice per group) are indicated in the figure panels. Statistical significance was determined by one-way ANOVA, with **p < 0.01, ***p < 0.001, and ****p < 0.0001.

Next, we tested whether β-catenin agonists function specifically through β-catenin pathways. WT mice injected with pristane (untreated with any drugs), WT mice injected with pristane and vehicle, and β-catenin cKO mice injected with pristane and treated with β-catenin agonists were compared. WT mice injected with pristane, either untreated or treated with vehicle, showed significant lung tissue damage (Fig. 7K–L, N-O & Q). Similarly, CAT-KO mice treated with β-catenin agonists also exhibited significant lung damage (Fig. 7M, P-Q). These results demonstrate that pharmacologic β-catenin agonists effectively mimic genetic stabilization, conferring protection against pristane-induced PH by suppressing pro-inflammatory cytokines, preserving organ integrity, and enhancing Treg-mediated repair pathways. Together, both genetic and pharmacologic activation of β-catenin establish a strong protective axis in PH, prompting us to next evaluate whether β-catenin agonists could suppress proinflammatory cytokines as shown in the genetic models.

### β-Catenin agonists reduce proteinuria and proinflammatory cytokines in the PH model

To evaluate whether β-catenin agonists suppress proinflammatory cytokines, we induced PH as described above (Fig. 7–8A). Urine analysis confirmed that WT mice injected with Pristane and treated with vehicle alone developed significantly higher proteinuria compared to pre-Pristane levels. In contrast, WT mice treated with β-catenin agonists showed no significant differences in proteinuria before and after Pristane injection (Fig. 8B–C).

**Figure 8.**
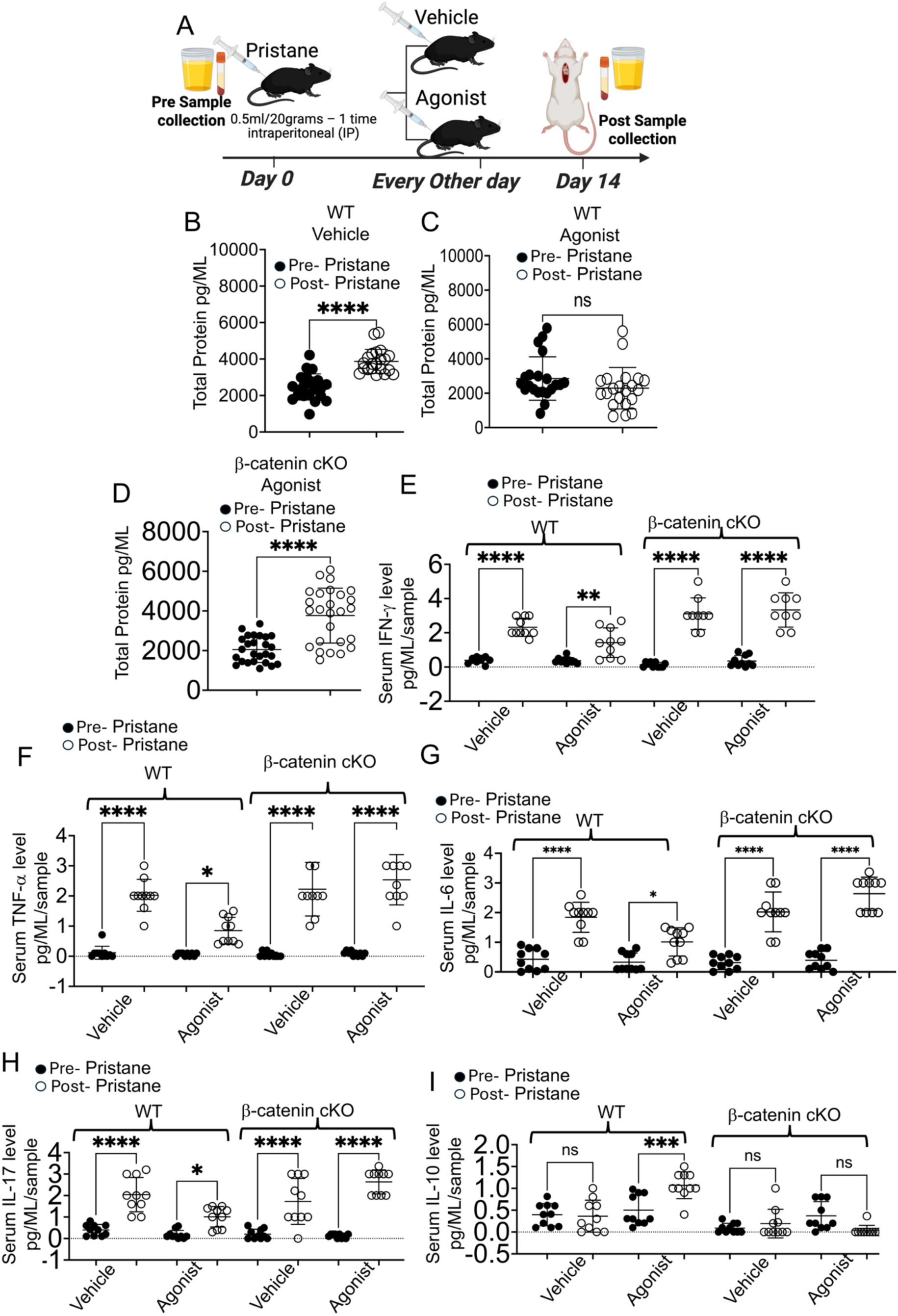
β-catenin agonists reduce proteinuria and pro-inflammatory cytokines in the PH model. (A) Schematic representation of the experimental design. Four groups of mice were used: two groups of WT mice and two group of β-catenin cKO mice. Urine and blood were collected via tail vein prior to Pristane injection. All groups received 0.5 ml/20 g Pristane. On day 14, mice were euthanized, and urine and serum were collected. (B–D) Proteinuria was measured by ELISA in each group, with quantitative comparison of pre- and post-Pristane samples. (E–H) Serum cytokines (IFN-γ, TNF-α, IL-6, IL-18, and IL-10) were quantified in WT mice treated with vehicle, WT mice treated with β-catenin agonist, and β-catenin cKO mice treated with vehicle or β-catenin agonist. Comparisons were made between pre- and post-Pristane samples. Quantitative data are presented as mean ± standard error of the mean (SEM). Sample sizes (n = 15–25 mice per group) are indicated in the figure panels. Statistical significance was determined by one-way ANOVA, with **p < 0.01, ***p < 0.001, and ****p < 0.0001.

Consistent with these findings, β-catenin agonist–treated mice exhibited significantly lower levels of IFN-γ and TNF-α compared to untreated or vehicle-treated controls. However, the reduction was incomplete and did not reach the degree of suppression observed in the genetic model or with Treg transfer (Fig. 8E–F). Similarly, IL-6 and IL-18 were reduced, although not to the same extent as in the genetic approaches (Fig. 8G–H). Notably, β-catenin agonist treatment significantly increased IL-10 production compared to vehicle-treated or untreated controls (Fig. 8J). In contrast, no significant differences were observed in IL-12 or IL-13 levels (Suppl. Fig. 4). Taken together, these findings demonstrate that β-catenin agonists enhance Treg responses in the lungs and suppress proinflammatory cytokines, highlighting their potential as a therapeutic intervention for PH.

### β-Catenin stabilization regulates gene expression during PH

Our data confirmed that β-catenin stabilization protects the lungs of our mice from PH. Further, these findings confirmed that β-catenin stabilization increases Tregs, which highly express Areg and BATF, and that these cells play a central role in tissue repair and reduction of inflammatory cytokines. However, the transcriptomic pathways regulated by β-catenin stabilization were still unknown.

To determine how β-catenin stabilization regulates transcriptomic pathways during pulmonary hemorrhage (PH), we induced PH with Pristane as described above. Animals were euthanized at days 14 and 21, and lungs tissues were collected and frozen for bulk RNA-sequencing. We examined differences in gene expression between WT and CAT-Tg post PH. Principal component analysis (PCA) of lung tissue identified two clusters of samples, which separated the WT and CAT-Tg groups (PC1: 53.05 %, PC2: 25.60% and PC3: 12.05%) (Fig. 9A). Further analysis of lung cell populations identified 2,688 differentially expressed genes (DEGs; FDR £ 0.05, |log2FC| ≥ 0.5) between WT and CAT-Tg (Fig. 9B), of which 1464 genes were downregulated, and 1224 genes were upregulated. DEGs and their normalized expression (row-scaled) were clustered by hierarchical clustering (Euclidean distance, complete linkage); the row dendrogram was cut into **k = 2** modules and module assignments were used to display expression clusters (Fig. 9C). Genes that were down and upregulated in CAT-Tg samples clustered in Module 1 and Module 2 respectively (Fig. 9C). GO enrichment analysis of Module 1 revealed that downregulated DEGs are involved in numerous biological pathways including response to stress, immune system process, protein binding, defense response and the inflammatory response. GO enrichment analysis of upregulated DEGs in Module 2 revealed genes involved in numerous biological pathways including cell projection assembly, plasma membrane bounded cell projection assembly, cell projection organization, and cell motility. GSEA analysis using the Hallmark pathways collection from Molecular Signatures Database (MSigDB)(94) revealed that the TNFα signaling via NF-κB, Inflammatory response, IL6–JAK–STAT3 signaling, mTORC1 signaling, MYC targets V1, E2F targets, G2M checkpoint, Epithelial–mesenchymal transition, Unfolded protein response, Hypoxia, MYC targets V2, KRAS signaling up, IL2–STAT5 signaling, p53 pathway, Apoptosis, Reactive oxygen species pathway, Interferon-γ response, Allograft rejection, UV response up, TGF-β signaling, Oxidative phosphorylation, Complement, Glycolysis, Interferon-α response, Androgen response, Mitotic spindle, Angiogenesis, Cholesterol homeostasis, DNA repair, PI3K–AKT–mTOR signaling, Coagulation, Estrogen response late, Protein secretion, Estrogen response early, and Heme metabolism pathways were enriched in WT compared to *Cat-Tg* in pre-transplanted CD8^+^ samples (Fig. 9D, 9E, 9F, 9G, 9H). KRAS signaling DN was enriched in CAT-Tg compared to WT.

**Figure 9.**
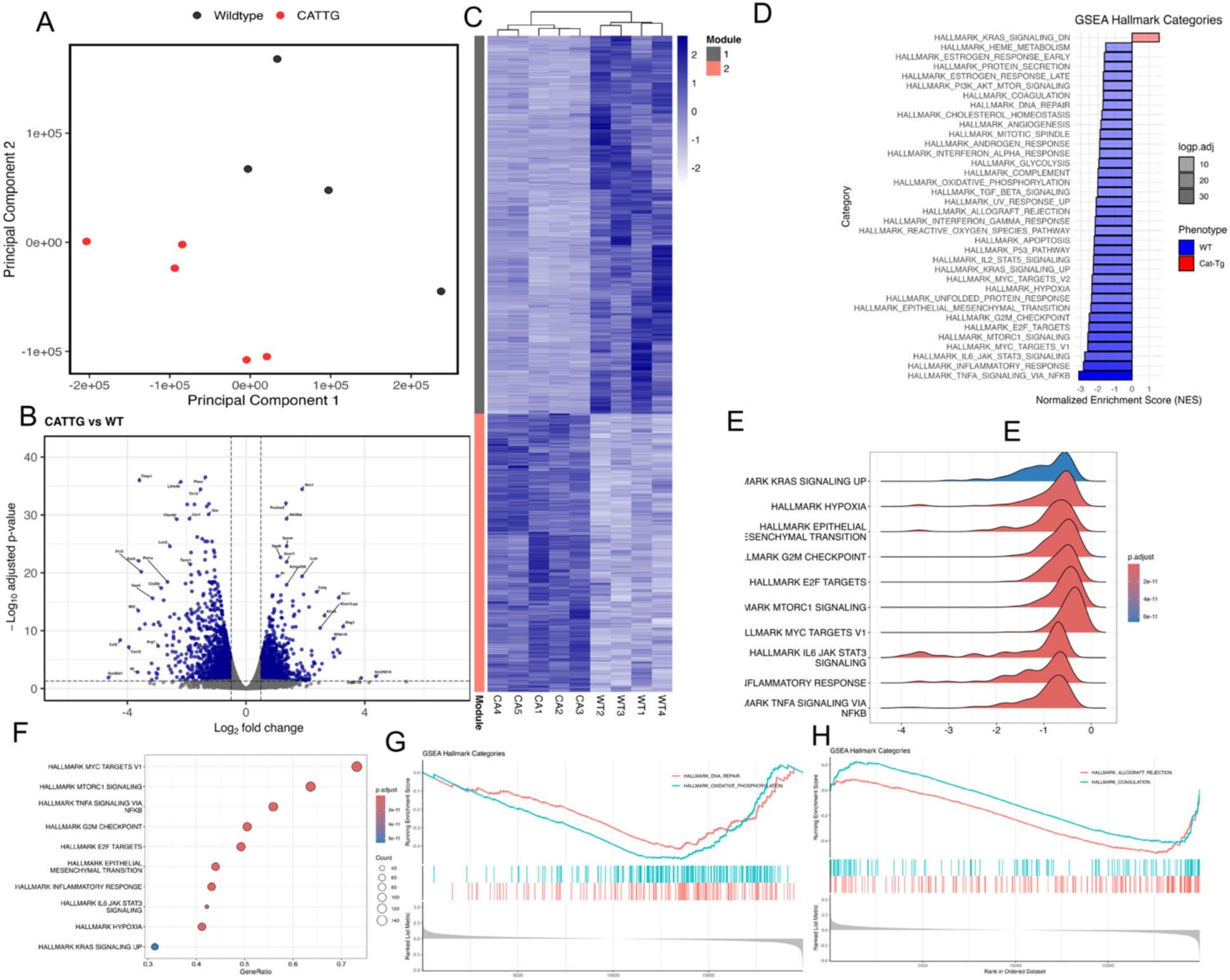
β-catenin overexpression differentially regulates gene expression in induced PH lung cells. (A) 2D graph of PCA analysis showing clustering of CAT-Tg and Wildtype lung cells by color. All replicates are shown (n = 4). (B) Volcano plot displaying 2,688 differentially expressed genes (DEGs; FDR ≤ 0.05, |log2FC| ≥ 0.5) between Cat-Tg and WT. Positive log2fold change corresponds to increased expression in Cat-Tg samples. (C) Hierarchical clustering and heat map illustrating expression of genes compared between different groups. All replicates are shown (n = 4) for each group. (D) A bar plot showing up or downregulated pathways based on enrichment scores in GSEA in WT or Cat-Tg in induced PH lung cell samples. Negative Normalized Enrichment Score (NES) is an indicator of downregulation and positive NES is an indicator of upregulation of the genes in CAT-Tg cell, in the corresponding pathway. Color specifies the group (WT or Cat-Tg) in which expression is altered (phenotype), color transparency indicates the negative Log10 of adjusted P-value. (E) A ridge plot showing the distribution of enrichments scores for selected Hallmark gene sets. Score (NES), is an indicator of downregulation and positive NES is an indicator of upregulation of the genes in CAT-Tg cell, in the corresponding pathway. Adjusted p-values were mapped to a continuous color scale from red (small adjusted p-values) to blue (large adjusted p-values). (F) A dot plot comparing the gene ratios for selected Hallmark gene sets. Size corresponds to count, the number of genes contributing to the enrichment. Adjusted p-values were mapped to a continuous color scale from red (small adjusted p-values) to blue (large adjusted p-values).(G–H) GSEA enrichment plots of selected gene clusters. (G) HALLMARK DNA REPAIR, HALLMARK OXIDATIVE PHOSPHORYLATION (H) HALLMARK ALLOGRAFT REJECTION, HALLMARK COAGULATION

These findings demonstrate that β-catenin stabilization reprograms lung transcriptomes during PH by suppressing inflammatory and stress-response pathways while promoting cell motility and tissue repair programs, thereby establishing a protective and regenerative environment.

### β-Catenin agonists impact gene expression during PH

Next, we wanted to determine whether pharmacological stabilization of β-catenin could mimic transcriptomic expression during PH. Thus, a cohort of WT mice were injected with Pristane as described above, then treated with vehicle or β-catenin agonists. At day 14, the mice were euthanized and the lungs were isolated from each mouse for bulk RNA sequencing. Our RNA seq analysis of the drug and vehicle treated cell populations identified 2565 differentially expressed genes **(DEGs; FDR** £ **0.05, |log2FC| ≥ 0.5)** between drug and vehicle samples **(Fig. 10B)**, of which 1583 genes were downregulated and 982 genes were upregulated. Principal component analysis (PCA) of drug and vehicle treated tissue identified two clusters of samples, identified as drug and vehicle groups (PC1: 65.84 %, PC2: 20.22% and PC3: 6.82%) (Fig. 9A). GO enrichment analysis of Module 1 revealed that downregulated DEGs are involved in the Profilin 1 complex, MCM complex, Trip(Br1)-Dp1-E2F1 complex, PLC-gamma-2-Lab-Blnk complex, BCR stimulation, immune system processes, immune responses, responses to stress, and defense responses. Go enrichment analysis of Module 2 revealed that upregulated DEGs are involved in cilium movement, microtubule bundle formation, microtubule-based movement, axoneme assembly, cilium-dependent cell motility, cilium or flagellum-dependent cell motility, cilium movement involved in cell motility, cilium organization, cilium assembly, microtubule-based process, flagellated sperm motility, sperm motility, cell projection assembly, and plasma membrane bounded cell projection assemblies. GSEA analysis showed that interferon gamma response, TNFα signaling via NF-κB, E2F targets, G2M checkpoint, mTORC1 signaling, inflammatory response, interferon alpha response, IL6–JAK–STAT3 signaling, epithelial–mesenchymal transition, KRAS signaling up, MYC targets V1, allograft rejection, complement, IL2–STAT5 signaling, hypoxia, p53 pathway, coagulation, apoptosis, unfolded protein response, glycolysis, mitotic spindle, MYC targets V2, angiogenesis, reactive oxygen species pathway, DNA repair, cholesterol homeostasis, UV response up and oxidative phosphorylation were negatively enriched in drug-treated tissue compared to vehicle-treated tissue in the harvested samples **(Fig. 10C, 10D, 10E, 10F, 10G).**

**Figure 10.**
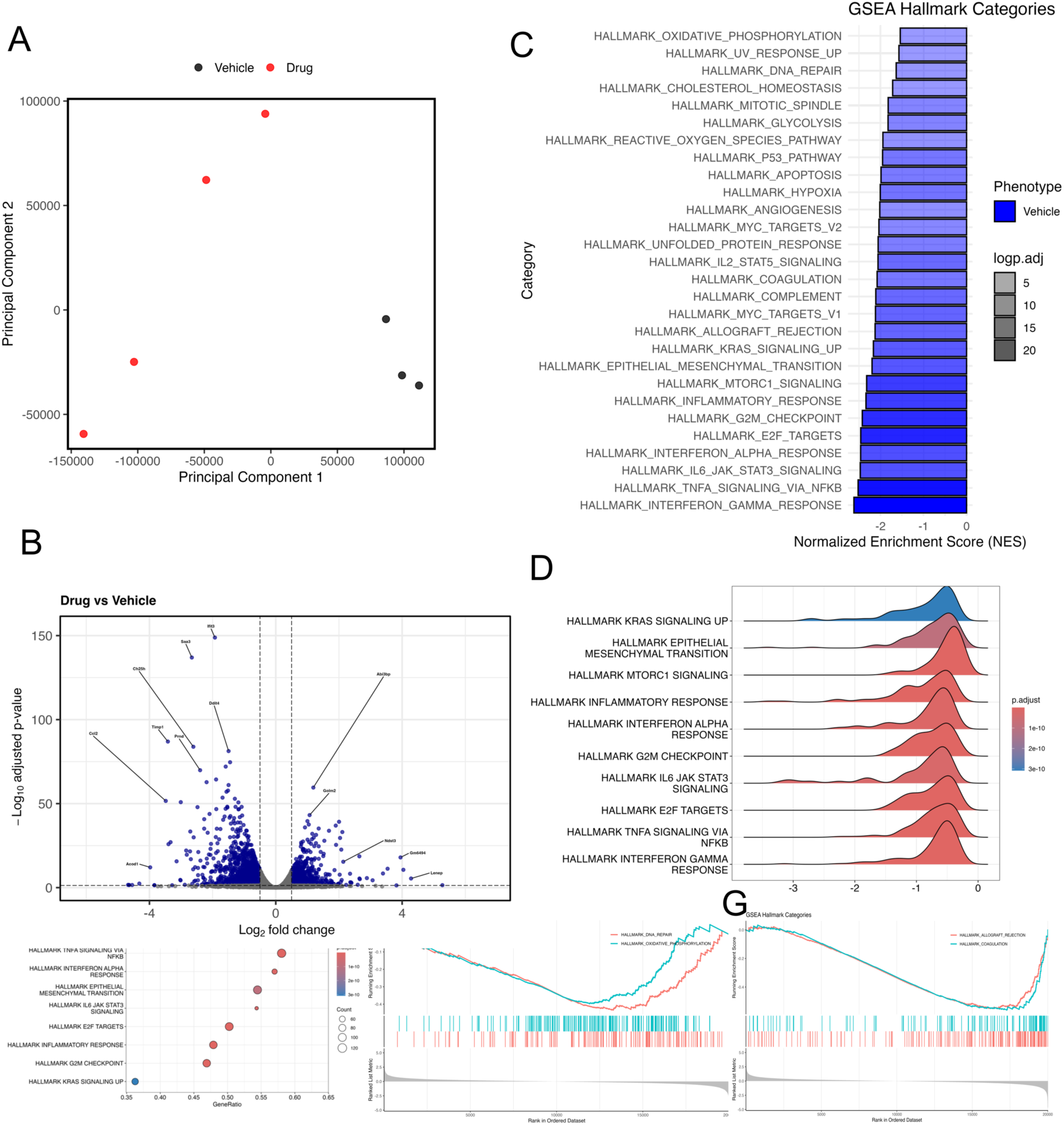
β-catenin overexpression differentially regulates gene expression in induced drug and vehicle treated cells. (A) 2D graph of PCA analysis showing clustering of drug and vehicle treated cells by color. All replicates are shown (n = 4, n = 3 for drug and vehicle respectively). (B) Volcano plot displaying 2,565 differentially expressed genes (DEGs; FDR ≤ 0.05, |log2FC| ≥ 0.5) between drug and vehicle. Positive log2fold change corresponds to increased expression in drug samples. (C) A bar plot showing up or downregulated pathways based on enrichment scores in GSEA in drug or vehicle in induced PH lung cell samples. Negative Normalized Enrichment Score (NES) is an indicator of downregulation and positive NES is an indicator of upregulation of the genes in drug cell, in the corresponding pathway. Color specifies the group (drug or vehicle) in which expression is altered (phenotype), color transparency indicates the negative Log10 of adjusted P-value. (D) A ridge plot showing the distribution of enrichments scores for selected Hallmark gene sets. Score (NES), is an indicator of downregulation and positive NES is an indicator of upregulation of the genes in drug treated cell, in the corresponding pathway. Adjusted p-values were mapped to a continuous color scale from red (small adjusted p-values) to blue (large adjusted p-values). (E) A dot plot comparing the gene ratios for selected Hallmark gene sets. Size corresponds to count, or the number of genes contributing to the enrichment. Adjusted p-values were mapped to a continuous color scale from red (small adjusted p-values) to blue (large adjusted p-values).(F–G) GSEA enrichment plots of selected gene clusters. (F) HALLMARK DNA REPAIR, HALLMARK OXIDATIVE PHOSPHORYLATION (G) HALLMARK ALLOGRAFT REJECTION, HALLMARK COAGULATION

Pharmacological β-catenin stabilization recapitulates the transcriptomic reprogramming seen in genetic models by suppressing inflammatory, stress-response, and proliferative signaling pathways while enhancing ciliary and motility-related programs, suggesting a therapeutic mechanism of protection during PH.

## Discussion

Pulmonary hemorrhage (PH) is defined as bleeding from the lower respiratory tract, manifesting clinically as hemoptysis or diffuse alveolar hemorrhage (DAH) (95-97). PH arises in the context of multiple conditions, including Goodpasture syndrome, systemic lupus erythematosus (SLE), and ANCA-associated vasculitis. In PH, autoimmune or inflammatory processes damage alveolar capillaries, leading to blood leakage into the alveolar spaces (1, 98-100). This intrapulmonary bleeding causes severe hypoxemia by impairing oxygen exchange, and recurrent or unresolved PH can drive long-term fibrosis and chronic respiratory disability (101-104).

Despite its clinical impact, the mechanisms by which regulatory T cells (Tregs) ameliorate PH remain incompletely understood. We recently reported that T cells from CAT-Tg mice can clear tumors without inducing graft-versus-host disease (GVHD) in an allogeneic transplant model (58). In that study, CAT-Tg T cells exhibited an activated phenotype, with high frequencies of central and effector memory subsets and increased expression of CD44, CD122, Eomes, and T-bet (Fig. 1–2). Consistent with this, we and others have shown that attenuated TCR signaling— whether via reduced TCR strength or decreased TCF-1 expression—promotes Treg generation (59, 84, 85). Here, we demonstrate for the first time that stabilizing β-catenin under the Lck promoter (43) recapitulates these effects, expanding Tregs in a manner similar to T cells with attenuated TCR signaling. β-Catenin stabilization was previously shown to enhance thymocyte development (43), our findings extend this paradigm by showing that β-catenin also promotes the activation of circulating T cells.

To directly test whether β-catenin stabilization protects against PH, we utilized a murine PH model. PH is a feature of several respiratory diseases, including bronchiectasis, tuberculosis, aspergilloma, cystic fibrosis, and tumors, where it frequently manifests as hemoptysis (7-10). Our data show that constitutive β-catenin expression preserves lung integrity in PH. Specifically, WT and β-catenin cKO mice exhibited significantly greater pulmonary damage than CAT-Tg mice, whereas no significant differences were observed in liver, kidney, or spleen pathology within the first 14 days. Notably, WT mice injected with pristane developed proteinuria early, and after 4–6 weeks, both kidneys and spleens displayed pathology (data not shown). These findings suggest that β-catenin’s protective effects are lung-specific in early disease but may influence systemic pathology over time.

Previous studies have consistently demonstrated that PH is associated with elevated proinflammatory cytokines (105-108). However, the mechanisms by which β-catenin stabilization counters this cytokine storm were unknown. Our results reveal that β-catenin stabilization reduces phosphorylation of STAT1, STAT3, and JAK1—key regulators of IFN-γ, IL-6, TNF-α, IL-18, IL-17, and IL-4 production (64, 84). Consistently, ELISA assays demonstrated that CAT-Tg mice exhibited reduced IFN-γ, IL-6, TNF-α, IL-18, IL-17, and IL-5 during PH, while anti-inflammatory cytokines such as IL-4, IL-13, and IL-10 were increased (109, 110). These data identify β-catenin as a regulator of cytokine storm severity, with therapeutic implications for PH and other cytokine-driven diseases.

Tregs are central to immune homeostasis and play a critical role in suppressing inflammation and preventing cytokine storms (59, 84, 111-113). However, induced Tregs often lose FOXP3 expression and stability in culture (114, 115), while natural Tregs exist in limited numbers. Importantly, our data show that CAT-Tg mice exhibit a significant increase in Tregs compared with WT mice, whereas loss of β-catenin markedly reduces Treg frequency. Moreover, CAT-Tg mice harbor substantially more Tregs in the lungs than WT controls (Fig. 5A–E). To investigate underlying mechanisms, we focused on Amphiregulin (AREG) and BATF, transcriptional regulators of tissue repair (20-26). β-catenin stabilization drives transcriptional programs including direct induction of AREG (27, 33). Consistent with this, CAT-Tg lungs expressed higher levels of AREG and were protected from PH(29, 34). Adoptive transfer studies confirmed that CAT-Tg–derived Tregs rescued recipients from PH, suppressing IFN-γ, TNF-α, and IL-17 while enhancing IL-10 production (Fig. 6). Together, these findings establish a mechanistic link between β-catenin stabilization, Treg expansion, and tissue repair during PH.

While genetic models provide strong mechanistic insights, they are not always directly translatable to the clinic. We therefore evaluated pharmacological approaches to β-catenin stabilization. WT mice treated with β-catenin agonists displayed increased Treg frequencies, reduced proinflammatory cytokines, and elevated IL-10 expression, resulting in significant protection against PH (Fig. 7). Our data demonstrate that pharmacological β-catenin activation recapitulates, to a large extent, the protective effects observed with genetic β-catenin stabilization. These findings provide a compelling rationale for therapeutic targeting of β-catenin in PH and potentially other cytokine storm–mediated diseases.

To understand how β-catenin stabilization impacts transcriptomic regulation during PH, our RNA-seq data demonstrate that β-catenin stabilization reprograms lung transcriptomes by suppressing inflammatory and stress-response pathways while promoting cell motility and tissue repair programs, thereby establishing a protective and regenerative environment. Furthermore, we show that pharmacological β-catenin stabilization closely recapitulates the transcriptomic reprogramming observed in genetic models by suppressing inflammatory, stress-response, and proliferative signaling pathways while enhancing ciliary and motility-associated programs, highlighting a therapeutic mechanism of protection during PH.

Together, these findings establish β-catenin stabilization—whether genetic or pharmacologic—as a central regulator of protective transcriptomic programs during PH, highlighting its therapeutic potential to suppress inflammation and promote tissue repair.

## Materials and Methods

### Mice

CAT-Tg mice were previously described (116) and were generously provided by Dr. Sen. β-catenin Flox/Flox mice were also a kind gift from Dr. Sen and were bred with CD4-Cre mice as described (57). C57BL/6, C57BL/6.SJL (B6-SJL), and B6-Ly5.1 (B6.SJL-Ptprca Pepcb/BoyCrl) mice were purchased from Charles River Laboratories or The Jackson Laboratory. Mice between 8 and 12 weeks of age were used in all experiments, with age- and sex-matched controls included. Sex as a biological variable, we used both male and female in each experiment.

All animal maintenance and experimental procedures were conducted in accordance with the guidelines of the Institutional Animal Care and Use Committee (IACUC) of SUNY Upstate Medical University (protocol #443).

### Reagents, cell lines, flow cytometry

Monoclonal antibodies were purchased from BioLegend or eBioscience and used at a 1:100 dilution unless otherwise specified. The following antibodies were used: anti-mouse CD3 (Cat# 100102), anti-CD28 (Cat# 102116), anti-CD3-BV605, anti-CD4-PE, anti-CD8-PE/Cy7, anti-Eomes-PE/Cy7, anti-CD44-Pacific Blue, anti-CD122-APC, anti-CD62L-APC/Cy7, anti-T-bet-BV421, anti-TNF-α-FITC, anti-IFNγ-APC, anti-CD45.1-FITC, anti-CD122-APC, anti-TCF-1-PE, and anti-CD45.2-APC. Additional antibodies included anti-FOXP3-Pacific Blue, anti-FOXP3-APC, anti-CD25-FITC, and anti-CD25-APC. Appropriate fluorochrome-matched isotype control antibodies (BioLegend/eBioscience) were used to establish gating thresholds. Flow cytometry was performed on a BD LSRFortessa (BD Biosciences). Dead cells were excluded using a live/dead viability dye (BioLegend). Singlets were identified based on forward and side scatter properties. CD4⁺ and CD8⁺ T cells were gated from live CD3⁺ T cells, and subsequent markers (e.g., CD44, CD62L, T-bet, Eomes, TCF-1, FOXP3) were analyzed within these subsets. Data were analyzed using FlowJo software (Tree Star, Ashland, OR) as shown in our publications (84, 85).

### Consumable

All Consumable were purchased from EIMMUNA Medical supply.

### Proteinuria Assays

Urine was collected day 0 pre pristane injection and day 14 or day 21 post injection. Using KIT protein BSA assay kit Pierce by scientific (23225). Pre-dilute urine was diluted 3 μl of urine to 27 μl of Phosphate-Buffer Saline (PBS). BCA (Bichinchonic Acid Assay) working reagent (WR) was prepared using 50:1 ratio of reagent A:B. 25μl of diluted urine was introduced to 200 μl of BCA WR in a 96 well V-bottom plate. The plate was then centrifuged for 30 seconds at 1250 rmp and shortly after incubated at ∼37 °C for 30 minutes. Samples were read using the Biotek Flx800 Fluorescene Microplate Reader at an absorbance was read at 562 nm. Proteinurina concentrations were analyzed using Gen5 and the stewarded curve was calculated for by comparison of BSA standards vs. sample concertation μg/mL.

### Cytokine production, assays

On day 14 pre and post treatment, serum was collected from cardiac blood. Pre- and Post-treatment Cytokine levels, including IFN-γ, TNF-α, IL-5, IL-12, IL-6, IL-10, IL-9, IL-17A, IL-17F, IL-22, and IL-13, were quantified using a multiplex bead-based immunoassay (LEGENDplex™, BioLegend) according to the manufacturer’s instructions (50, 59). Samples were acquired on a BD LSRFortessa flow cytometer (BD Biosciences), and cytokine concentrations were analyzed with LEGENDplex Data Analysis Software (BioLegend).

### Western blotting

Cells were lysed in freshly prepared RIPA buffer (Fisher Scientific, Cat# PI89900) supplemented with complete protease inhibitor cocktail (Sigma-Aldrich, Cat# 11697498001) and centrifuged at 14,000 rpm for 10 minutes at 4 °C. Aliquots containing 1 × 10X^6^ cells were separated on 12–18% SDS–polyacrylamide gels and transferred onto nitrocellulose membranes for immunoblotting. All cell culture reagents and chemicals were obtained from Invitrogen (Grand Island, NY) or Sigma-Aldrich (St. Louis, MO), unless otherwise specified. For signaling analysis, the following antibodies were used: anti-pSTAT1 (Cell Signaling Technology, Cat# 9167), anti-STAT1 (Cell Signaling Technology, Cat# 9172), anti-pSTAT3 (Cell Signaling Technology, Cat# 9131), anti-JAK1 (Cell Signaling Technology, Cat# 3332), anti-pJAK3 (Cell Signaling Technology, Cat# 5031), and anti-β-Actin (Cell Signaling Technology, Cat# 4970). Additional antibodies included Amphiregulin (AREG, clone AREG559, eBioscience/Thermo Fisher, Cat# 14-9999-82) and BATF (mouse monoclonal, Thermo Fisher, Cat# MA5). For immunohistochemistry, the following antibodies were used: anti-CD4 (clone GK1.5, Thermo Fisher, Cat# 14-0041-82), anti-CD45.2 (clone 104, Thermo Fisher, Cat# 14-0454-82), and anti-FOXP3 (clone FJK-16s, Thermo Fisher, Cat# 14-5773-82).

### Histopathological Evaluation

Mice were injected with Pristane and euthanized on day 14 or day 21 post-treatment. Lungs, liver, kidneys, and small intestine were collected and fixed in 10% neutral buffered formalin, followed by paraffin embedding, sectioning, and hematoxylin and eosin (H&E) staining by the Histology Core Facility at SUNY Upstate Medical University. Histopathological evaluation of pulmonary hemorrhage (PH) was performed by a board-certified pathologist (L.C.), who was blinded to both study groups and disease status. Tissue grading was conducted according to established criteria (117, 118). Statistical analysis of histopathological scores was performed using the Mann–Whitney U test.

### Regulatory T cells (Tregs) isolation

CAT-Tg and WT mice were bred onto the C57BL/6 background with the Foxp3 ^tm1^Flv/J strain, an X-linked targeted knock-in model in which Foxp3-expressing cells are co-marked with monomeric red fluorescent protein (mRFP). RFP expression faithfully reflects Foxp3 gene expression in lymphocytes. This strain was kindly provided by Dr. Avery August (Cornell University). We confirmed that both WT and CAT-Tg mice expressed FoxP3-RFP. CD4⁺ lymphocytes were isolated from spleens of CAT-Tg and WT mice using anti-CD4 magnetic microbeads and column-based separation (Miltenyi Biotech, Auburn, CA). Treg cells were identified by surface expression of CD3⁺, CD25⁺, and intracellular Foxp3⁺ (RFP⁺). Cells were subsequently sorted on a BD FACSAria IIIu cell sorter (BD Biosciences). The purity of sorted Treg populations consistently exceeded 98% unless otherwise specified. All cell culture reagents and chemicals were purchased from Sigma-Aldrich (St. Louis, MO) and Invitrogen (Grand Island, NY), unless otherwise noted as shown by our publications (51, 85).

### Statistics

All numerical data are reported as mean ± standard deviation (SD) unless otherwise indicated in the figure legends. Statistical analyses were performed using GraphPad Prism v10 (GraphPad Software, San Diego, CA). Group differences were evaluated using one-way or two-way ANOVA followed by Tukey’s multiple comparisons test, chi-square test, or unpaired Student’s *t*-test, as appropriate. A *p*-value ≤ 0.05 was considered statistically significant. For transplant experiments, sample size was set at *n* = 3 mice per group, and all experiments were independently repeated at least twice. Sample sizes were determined based on power analyses unless otherwise specified. Mice were sex- and age-matched as closely as possible, consistent with our prior publications (38, 48-51, 57-59, 84, 85, 119-129)

### RNA sequencing

To investigate the molecular basis by which β-catenin stabilization alters transcriptomic profiles, we employed two approaches. First, four WT and four CAT-Tg mice were injected with Pristane and euthanized on day 14; lungs from each mouse were collected and frozen. In the second approach, WT mice were injected with Pristane and subsequently treated with a β-catenin agonist. Tissues from both experiments were submitted to the Molecular Analysis Core Facility at SUNY Upstate Medical University for RNA extraction and library preparation, followed by RNA sequencing. RNA-seq data were generated from four groups: 1) WT- pre-treatment, 2) WT-post-treatment and CAT-Tg 3) pre-and 4) post treatment with pristane Data processing and analysis were performed using R (version 4.5.1) with the RStudio interface (version 2025.05.1+513) and Bioconductor packages. Transcript abundance was quantified by pseudoalignment with Kallisto (version v0.51.1) (130) Transcript-per-million (TPM) values were normalized for each sample simultaneously and fitted to a linear model using the **sleuth** R package (131, 132). Differentially expressed genes (DEGs) were defined as those exhibiting a log2fold change ≥ |.5| with a false discovery rate (FDR) ≤ 5%, after multiple testing correction using the Benjamini–Hochberg method (133). This data was used for hierarchical clustering and heatmap generation in R.

For pathway analysis, Gene Ontology (GO) enrichment was assessed with the **gprofiler2** R package (134) using the gost function. Gene Set Enrichment Analysis (GSEA) was performed with the **clusterProfiler** R package(135), and the Molecular Signatures Database (MSigDB) (94), focusing on the Hallmark pathways collection. RNA-seq data will be deposited in the NCBI Gene Expression Omnibus (GEO) database (https://www.ncbi.nlm.nih.gov/geo/ As shown in our publications (50-52, 58, 59).

## Author contributions

FM, HX, AM, MD, SM, RT, MM, and TD performed experiments LC, analysis histological analysis JMS, Provided CAT-Tg and β-catenin cKO mice. MC provided valuable reagents. MK designed experiments, analyzed the data, and wrote the manuscript provided reagents.

## Acknowledgements

We thank all members of the Karimi for helpful discussions. Supplemental material

---

**Summary Figure:**
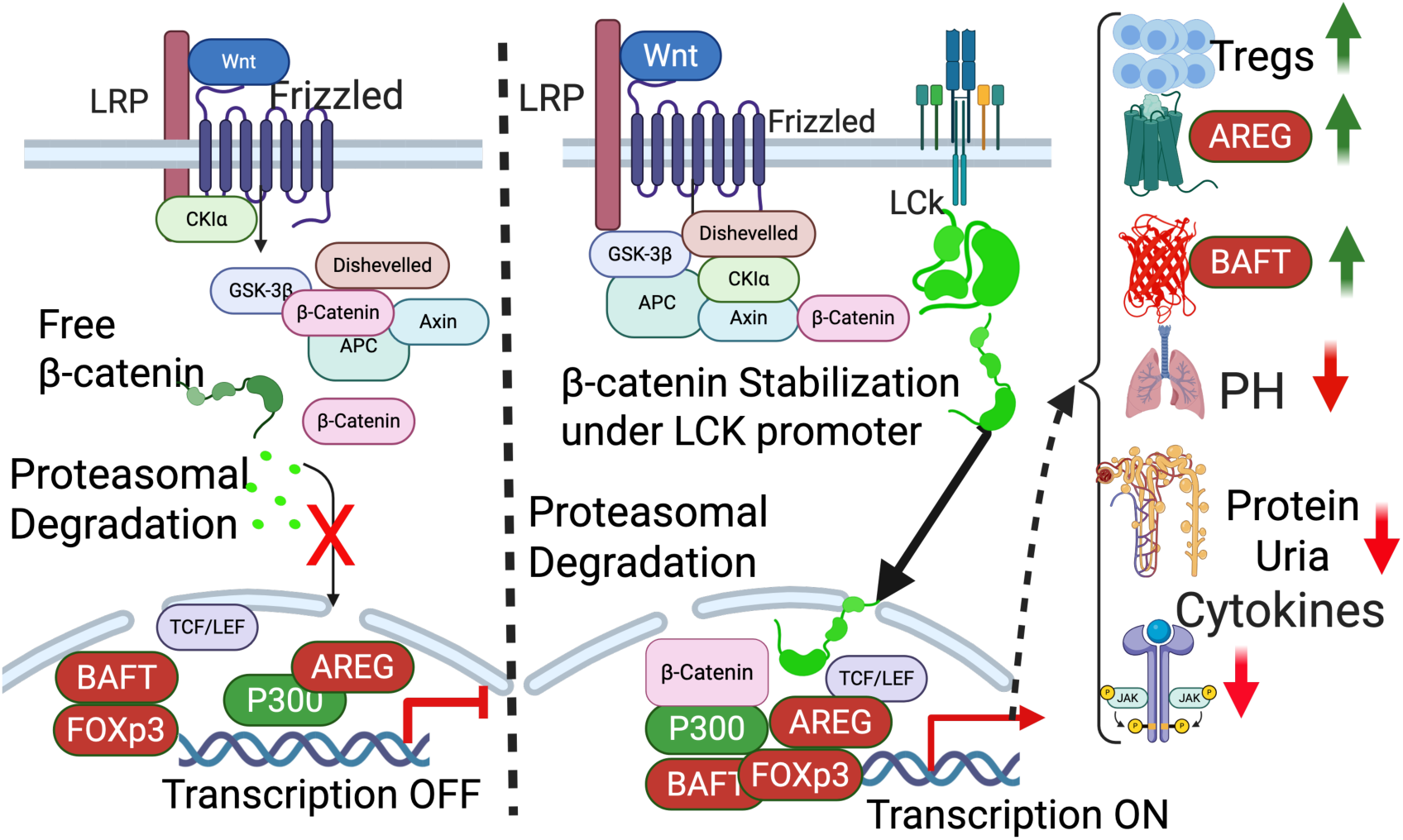
When a Wnt ligand binds to its receptors (Frizzled and LRP5/6), it triggers a cascade that leads to the stabilization and nuclear translocation of β-catenin. In the absence of Wnt signaling, β-catenin is targeted for degradation by a protein complex including GSK-3β, CK1α, and APC. When β-catenin is degraded its does enter nucleus and transcription is off. CAT-Tg mice where β-catenin is stabilized its protected from proteasomal degradation, stabilizing it in the cytoplasm and safely transferring it to the nucleus and transcription is on where it functions as a transcriptional coactivator for TCF/Lef, P300, AREG, FOXP3, BAFT and Groucho. In CAT-Tg mice we observed increased in Tregs that have higher amount of BAFT and ARCH. That alleviate PH, decrease proteinuria and cytokine

**Supplementary Figure 1.**
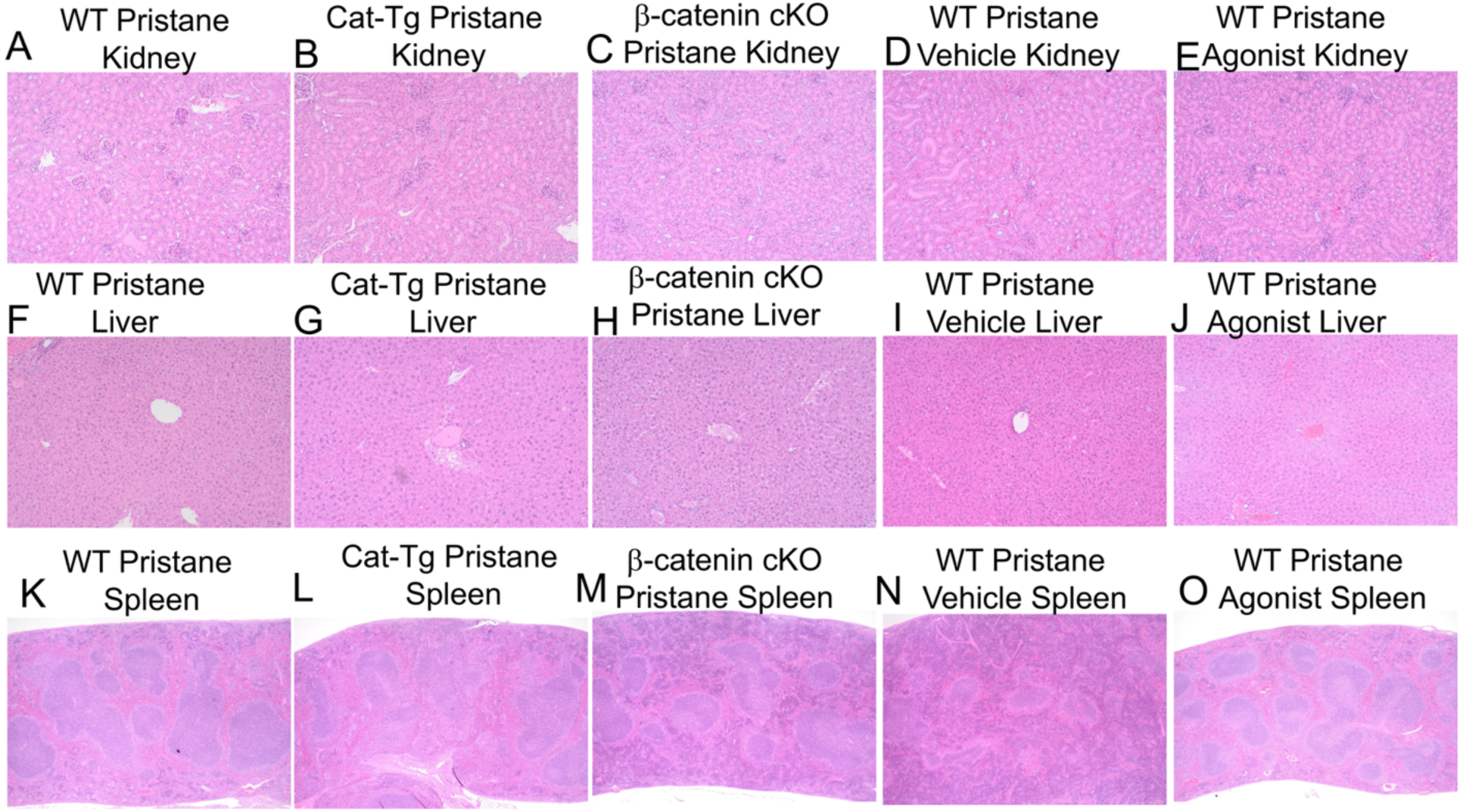
Short-term Pristane injection does not affect kidneys, liver, or spleen. (A–C) WT mice treated with Pristane were examined by histological analysis on day 14, showing no evidence of pulmonary hemorrhage (PH) or organ damage. (D–F) CAT-Tg mice treated with Pristane on day 14 similarly showed no histological signs of PH or damage. (G–I) β-catenin cKO mice treated with Pristane on day 14 showed no evidence of PH or damage. (J–L) WT mice treated with Pristane and rescued with vehicle alone displayed no histological abnormalities on day 14. (M–O) WT mice treated with Pristane and rescued with β-catenin agonist alone also showed no histological signs of PH or organ damage at day 14.

**Supplementary Figure 2.**
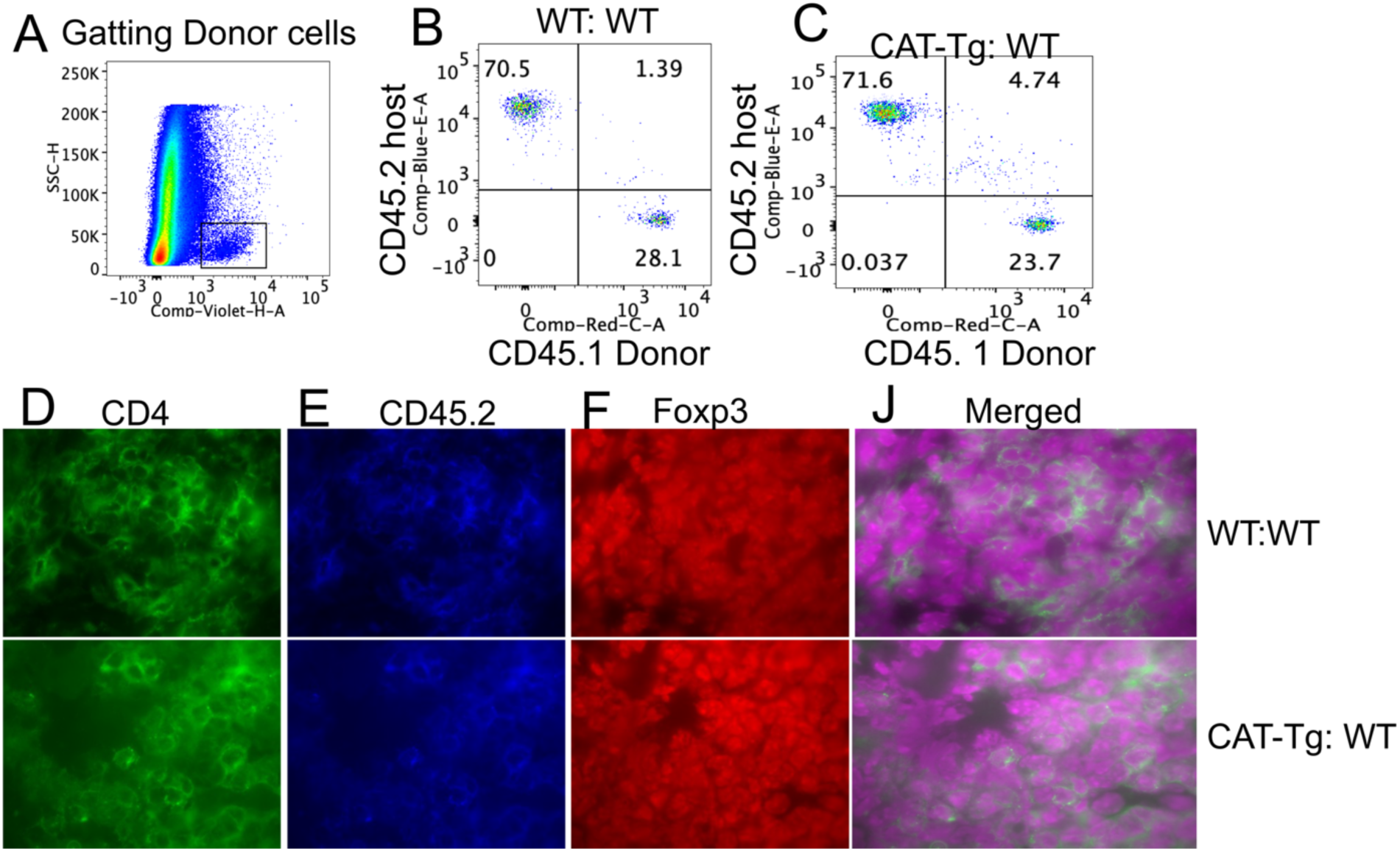
Presence of donor Tregs in host mice. (A–C) Donor Tregs expressing the CD45.2 congenic marker, isolated from WT or CAT-Tg mice, were transplanted into WT recipient mice carrying the CD45.1 congenic marker. Both donor and recipient mice were on a C57BL/6J background. On day 14, host mice were euthanized, and CD3⁺ T cells were analyzed. CD3⁺ cells were further subdivided into CD45.2⁺ donor-derived T cells and CD45.1⁺ host-derived T cells in the spleen. (D–J) To confirm the presence of donor Tregs in the lungs, immunohistochemistry was performed. Cells were stained for CD4 and CD45.2 (donor Tregs, red) and FOXP3 (green), with merged images showing co-localization of donor T cells and FOXP3 expression (blue nuclear counterstain).

**Supplementary Figure 3.**
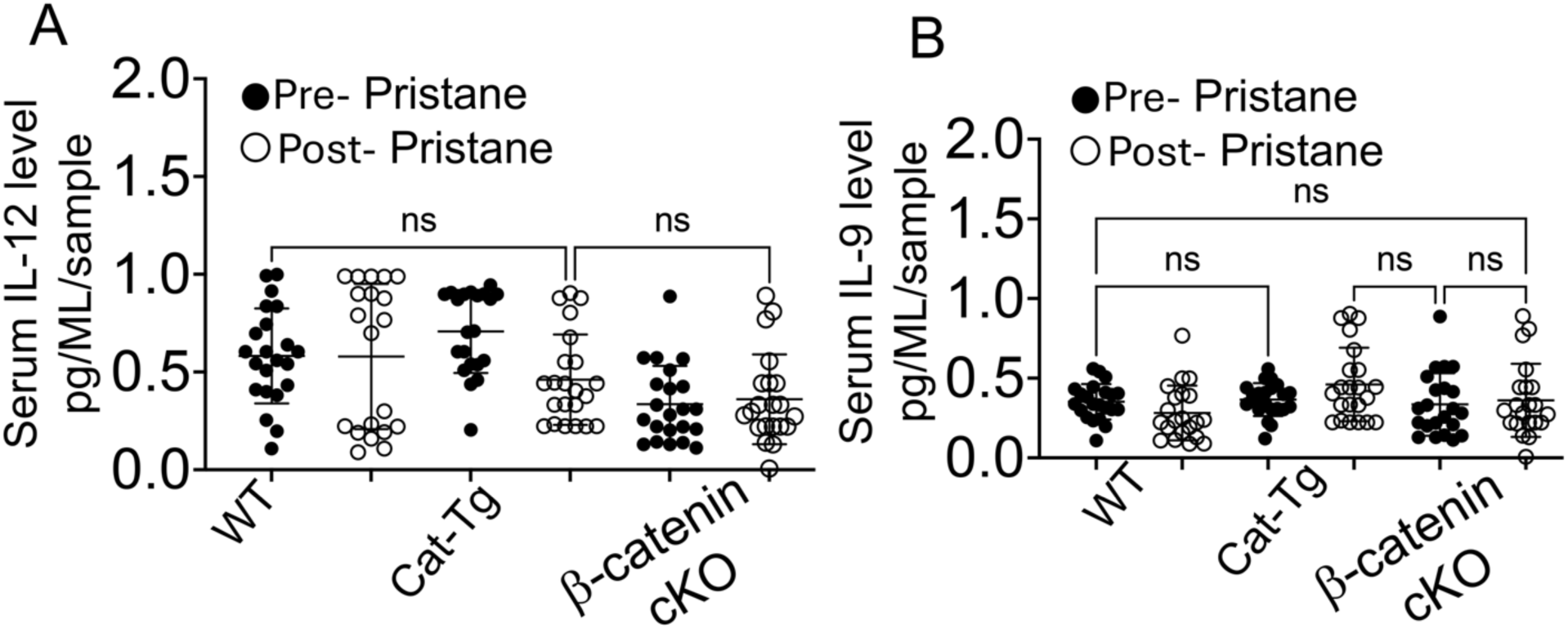
β-catenin stabilization does not impact IL-12 and IL-9 cytokines during PH. (A–B) Serum levels of IL-12 and IL-9 were measured by ELISA in WT, CAT-Tg, and β-catenin cKO mice before and after Pristane injection. Quantitative data are presented as mean ± standard error of the mean (SEM). Sample sizes (n = 15–25 mice per group) are indicated in the figure panels. Statistical significance was determined by one-way ANOVA, with **p < 0.01, ***p < 0.001, ****p < 0.0001, and NS indicating non-significant differences.

**Supplementary Figure 4.**
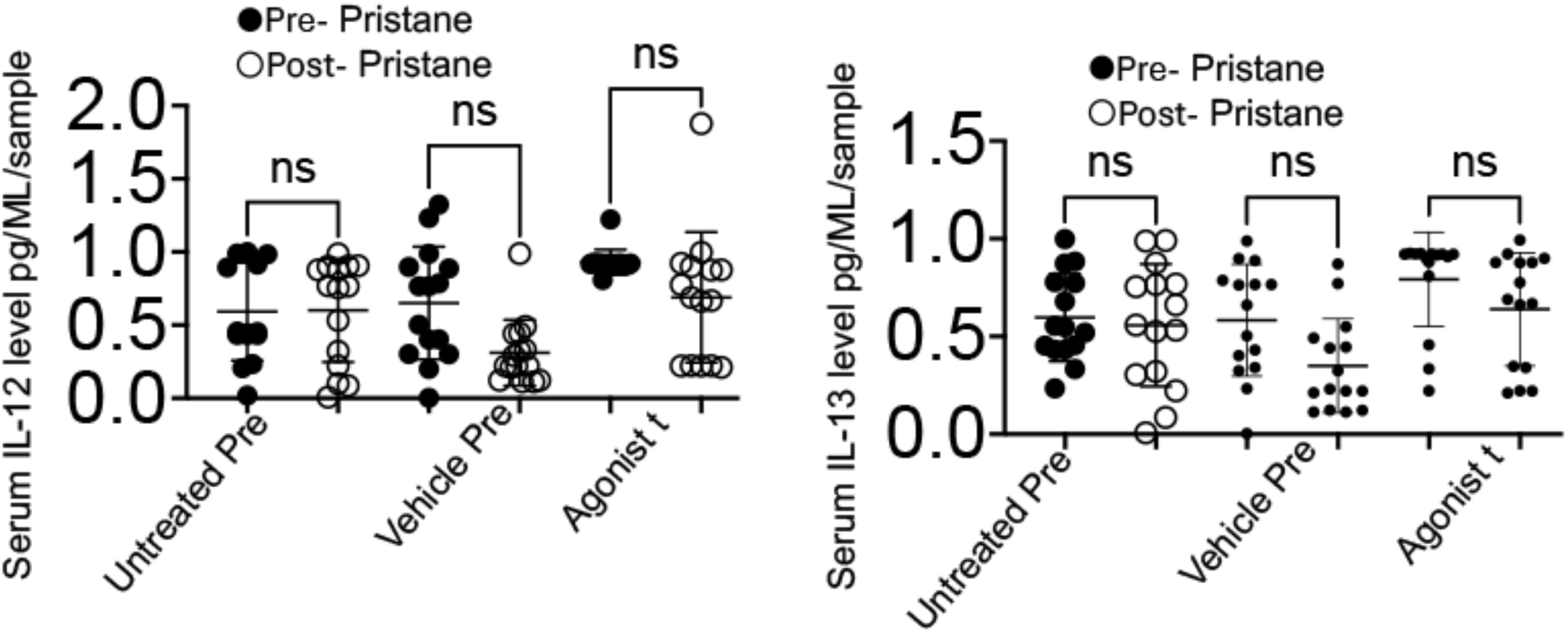
β-catenin agonists do not impact IL-12 and IL-13 cytokines during PH. (A–B) Serum levels of IL-12 and IL-13 were measured by ELISA in WT, CAT-Tg, and β-catenin cKO mice before and after Pristane injection. Quantitative data are presented as mean ± standard error of the mean (SEM). Sample sizes (n = 15–25 mice per group) are indicated in the figure panels. Statistical significance was determined by one-way ANOVA, with **p < 0.01, ***p < 0.001, ****p < 0.0001, and NS indicating non-significant differences.

