## Supplementary figures and images for "β-Catenin Stabilization Protects Against Pulmonary Hemorrhage Through Amphiregulin and BATF- Mediated Regulatory T Cells"

### Supp.Fig 1

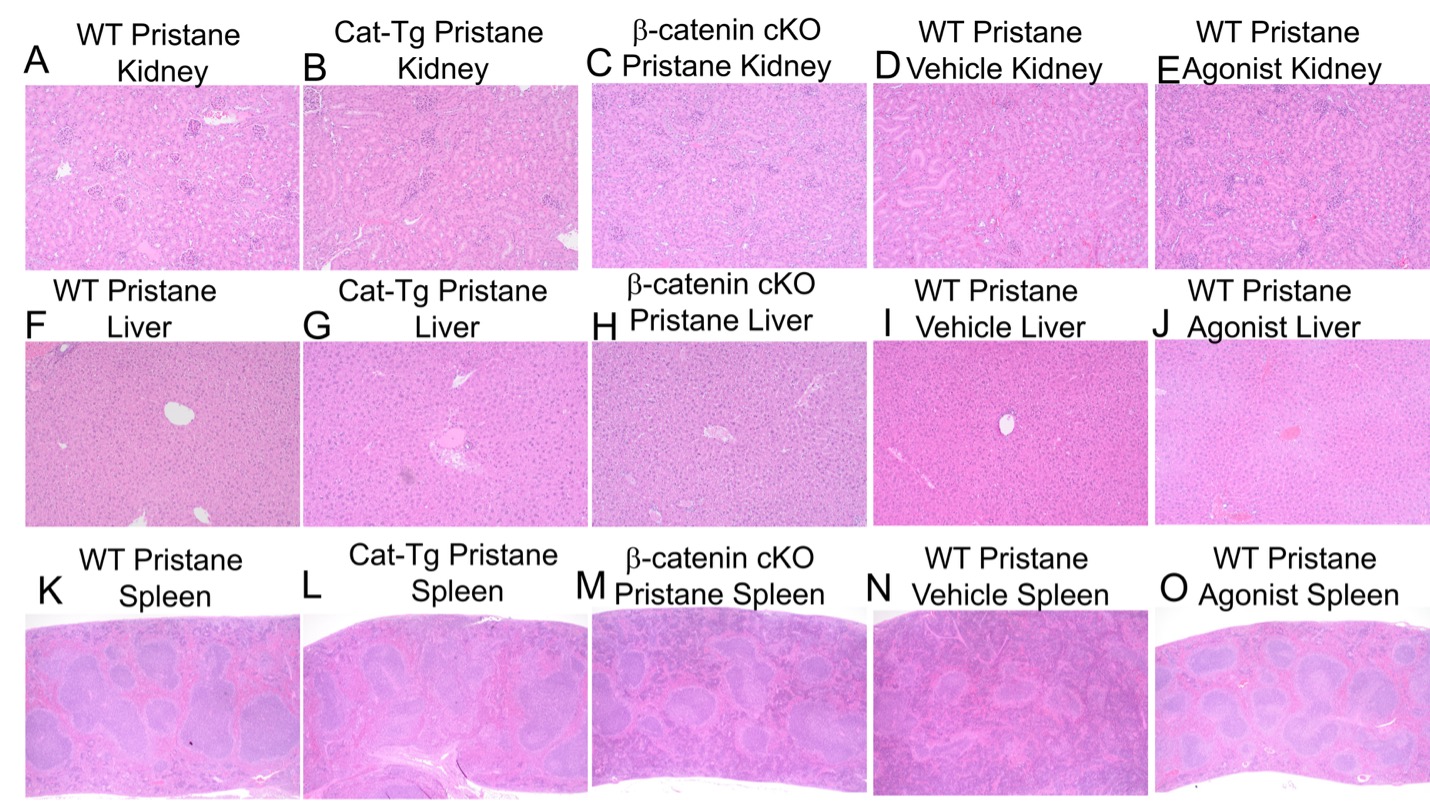

### Supp.Fig 2

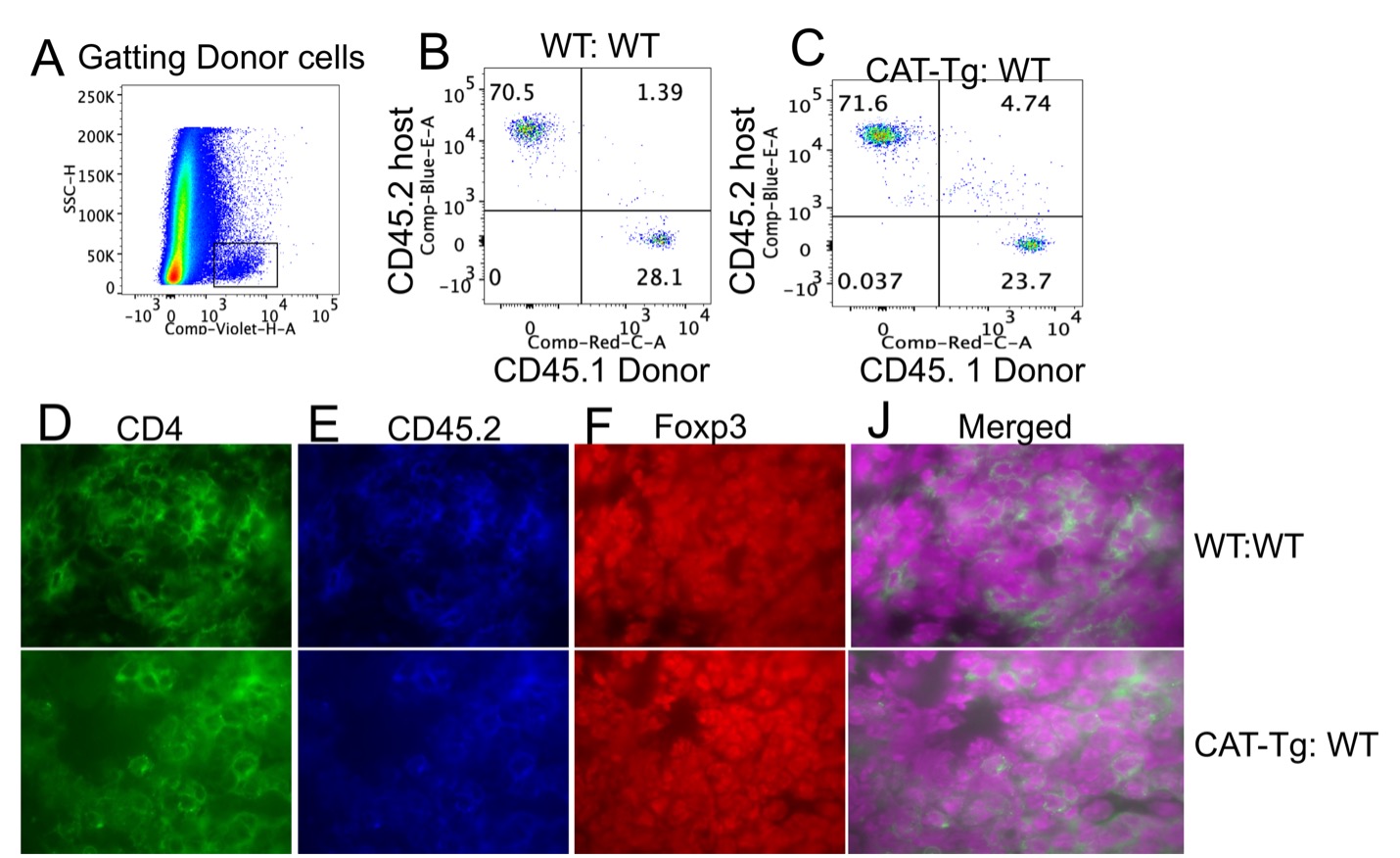

### Supp.Fig 3

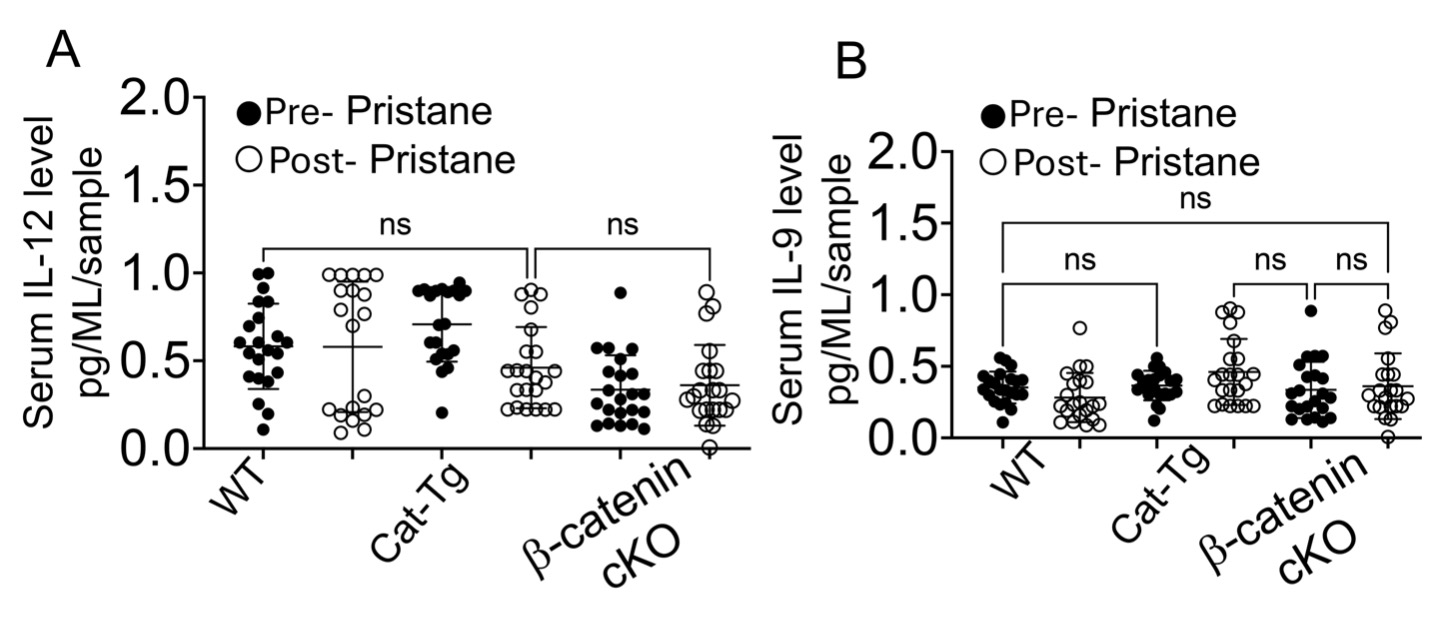

### Supp.Fig 4

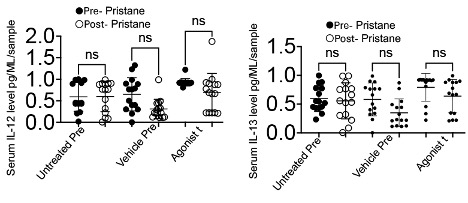
